# RNase L-induced bodies sequester subgenomic flavivirus RNAs and re-establish host RNA decay

**DOI:** 10.1101/2024.03.25.586660

**Authors:** J. Monty Watkins, James M. Burke

**Affiliations:** Department of Molecular Medicine, The Herbert Wertheim University of Florida Scripps Institute for Biomedical Innovation and Technology, Jupiter, FL, United States of America; Department of Immunology and Microbiology, The Herbert Wertheim University of Florida Scripps Institute for Biomedical Innovation and Technology, Jupiter, FL, United States of America; Skaggs Graduate School of Chemical and Biological Sciences, The Scripps Research Institute, Jupiter, FL, USA

**Keywords:** Flavivirus, sfRNA, RNase L-induced body, condensates, stress granule, P-body, RNA decay, RNase L

## Abstract

Subgenomic flavivirus RNAs (sfRNAs) are structured RNA elements encoded in the 3’-UTR of flaviviruses that promote viral infection by inhibiting cellular RNA decay machinery. Herein, we analyze the production of sfRNAs using single-molecule RNA fluorescence *in situ* hybridization (smRNA-FISH) and super-resolution microscopy during West Nile virus, Zika virus, or Dengue virus serotype 2 infection. We show that sfRNAs are initially localized diffusely in the cytosol or in processing bodies (P-bodies). However, upon activation of the host antiviral endoribonuclease, Ribonuclease L (RNase L), nearly all sfRNAs re-localize to antiviral biological condensates known as RNase L-induced bodies (RLBs). RLB-mediated sequestration of sfRNAs reduces sfRNA association with RNA decay machinery in P-bodies, which coincides with increased viral RNA decay. These findings establish a role of RLBs in promoting viral RNA decay, demonstrating the complex host-pathogen interactions at the level of RNA decay and biological condensation.

**Highlights:**

- Single-molecule imaging of sfRNA production and localization
- sfRNAs localize to RNase L-induced bodies
- RNase L-induced bodies sequester sfRNAs away from P-bodies
- Sequestration of sfRNAs by RNase L-induced bodies enhances decay of viral genomes

## INTRODUCTION

Dengue virus, West Nile virus, and Zika virus are mosquito-borne flaviviruses that cause severe and fatal diseases in humans, including hemorrhagic fever, microcephaly, and encephalitis.^1–3^ Currently, the territory of many flaviviruses is expanding,^4–7^ eradication of flaviviruses is difficult due to their mosquito reservoir, and vaccines approved for humans exist for only a fraction of flaviviruses.^8^ It will be imperative to develop treatments that reduce viral-associated pathogenesis, which will require a better understand the host-pathogen interactions between mammalian cells and flaviviruses.

Flavivirus genomes are positive-sense, single-stranded RNAs (+ssRNA) ∼11 kb in length that encode 5’- and 3’- untranslated regions (UTRs) and an open-reading frame encoding a polyprotein. During infection in both mosquitoes and mammalian cells, the structured 3’ UTRs are generated as an independent fragment in large quantities, and thus are termed subgenomic flavivirus RNAs (sfRNAs) ^9^. sfRNAs are generated when exoribonuclease 1 (XRN1), which is the primary 5’->3’ cellular RNA decay exoribonuclease that degrades decapped mRNAs,^10–15^ stalls on a dumbbell hairpin structural element that is conserved across the genus flavivirus during 5’-3’ decay of the flavivirus genome.^16–19^

The sfRNA structures contribute to cytopathogenic effects in culture mammalian cells and pathogenicity in mouse models.^9^ However, the functions of sfRNAs are not fully understood. In both insects and mammals, sfRNAs stall XRN1, which leads to an increase in uncapped cellular mRNAs.^17^ Thus, the stalling of XRN1 on sfRNAs is thought to sequester XRN1 to reduce decay of viral RNA. In addition, sfRNAs have been shown to inhibit antiviral responses, including the type I interferon response in mammals and RNA interference (RNAi) in mosquitoes.^20–22^

Ribonuclease L (RNase L) is a mammalian endonuclease that is activated upon the recognition of cytoplasmic viral double-stranded RNA (dsRNA) by oligoadenylate synthetase (OAS) proteins.^23^ OAS binding to dsRNA stimulates its production of oligo-2-5A. Oligo-2-5A binds RNase L and promotes homodimerization of RNase L,^23^ which activates the RNase domain of RNase L.^24^ Once activated, RNase L cleaves host and viral ssRNA at UN^N motifs.^24–26^ RNase L has the capacity to rapidly degrade nearly all cellular mRNAs following activation.^27,28^

Concurrent to the initiation of cellular mRNA decay mediated by RNase L, cells assemble cytoplasmic biological condensates known as RNase L-induced bodies (RLBs).^27^ RLBs concentrate several RNA-binding proteins, including PABPC1, G3BP1, and UBAP2L.^29,30^ Notably, many of these RNA-binding proteins can also localize to stress granules (SGs), which are large RNP complexes composed of non-translating mRNPs that form when translation initiation is repressed by phosphorylation of eIF2α.^31,32^ However, RLBs are distinct from SGs based on several criteria. First, unlike stress granules, RLBs do not require protein kinase R (PKR)-mediated phosphorylation of eIF2α or G3BP1-mediated RNA condensation for their assembly. Second, RLBs have a distinct proteome in comparison to SGs, and thus lack several RNA-binding proteins that typically localize to SGs, such as TIA-1. Third, RLBs stably associate with processing bodies (P-bodies),^29^ whereas SG association with P-bodies is transient.^33^ Lastly, unlike SGs, RLBs typically lack intact mRNAs but concentrate poly(A)+ RNA, suggesting that they enrich for 3’-end decay fragments of mRNAs.^29^

Herein, we use single-molecule RNA super-resolution microscopy to demonstrate that sfRNAs and larger 3’-end degradation fragments of flavivirus localize to RLBs in mammalian cells during Zika virus, West Nile virus, or Dengue virus serotype 2 infection. We show that RLB assembly reduces sfRNA localization to P-bodies, and that this coincides with robust XRN1-mediated decay activity and degradation of viral RNAs. These findings transform our understanding of sfRNA biology in mammalian cells and demonstrate that RLBs function to sequester pathogenic viral RNAs to enhance viral RNA decay.

## RESULTS

### Single-molecule RNA visualization of sfRNA production during virus infection

To examine sfRNA production and localization during flavivirus infection, we generated single-molecule RNA fluorescence *in situ* hybridization (smRNA-FISH) probe sets that target the sfRNA-encoding regions of dengue virus serotype II (DENV2), West Nile virus (WNV), or Zika virus (ZIKV) (**Fig. 1A**). To differentiate sfRNAs from the full-length genomes or longer 3’- degradation fragments, we generated probe sets specific to the first 1-kb (5’-CDS) or the last 1-kb (3’-CDS) of each virus, respectively (**Fig. 1A**). Each probe set (5’-CDS, 3’-CDS, sfRNA) was labeled with distinct fluorophores (ATTO-488, ATTO-550, ATTO-633, respectively) to allow for simultaneous imaging (**Fig. 1A**).

**Fig. 1.**
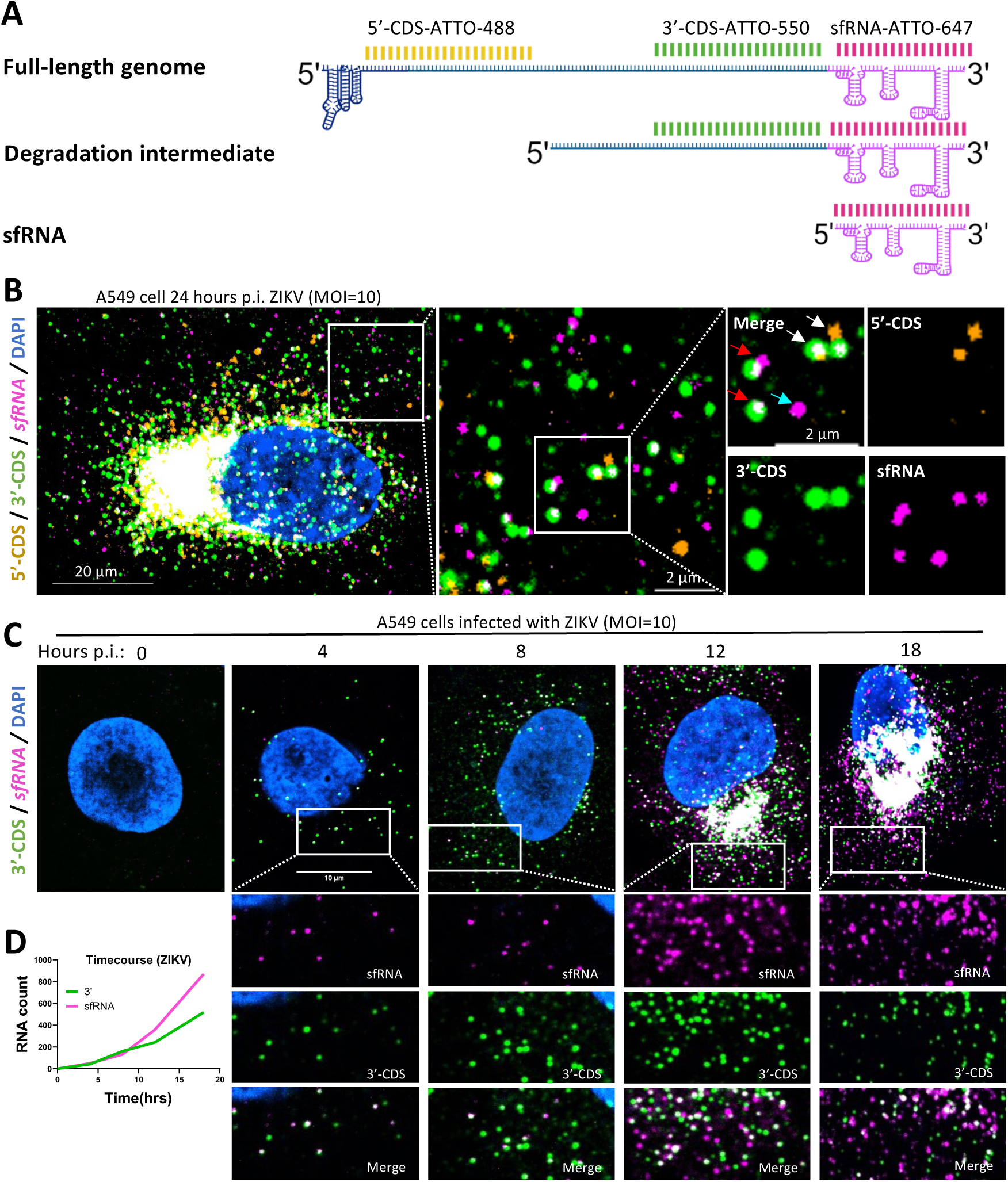
sfRNAs diffusely localize in the cytosol during early infection. (**A**) Schematic showing regions of the flavivirus genome and target locations of smRNA-FISH probes (not to scale) and examples of different RNA decay species. (**B**) Co-staining smRNA-FISH for ZIKV 5’-CDS, 3’-CDS, and sfRNAs 12 hours post-infection (p.i.) of A549 cells with ZIKV (MOI=10). Inset: The white arrow points to full-length genome, red to 3’ degradation fragment, and cyan to sfRNA. **(C**) Representative images of 3’-CDS and sfRNA staining ranging from pre-infection to 18 hours p.i. **(D)** Quantitation of average RNA count per cell from (C).

We confirmed that each probe set (5’-CDS, 3’-CDS, sfRNA) targeting DENV2, ZIKV, or WNV specifically detected viral RNA by staining cells with each probe set individually (**Fig. S1A**). While the probe sets (5’-CDS, 3’-CDS, sfRNA) did not stain mock-infected cells (**Fig. S1A**), each probe set (5’-CDS, 3’-CDS, sfRNA) intensely stained a region proximal to the nucleus 24 hours post-infection (p.i.) with DENV2, ZIKV, or WNV (**Fig. S1A,** yellow arrows). This structure is consistent with the morphology of the viral replication organelle (RO), which would be expected to concentrate full-length genomes and viral double-stranded RNA (dsRNA), which we confirmed by immunofluorescence assays for dsRNA (**Fig. S1B**). These data demonstrate that each probe set (5’-CDS, 3’-CDS, sfRNA) reliably detects the specific region of interest in each viral RNA.

We next simultaneously stained cells with all three probe sets 12 hours post-infection (p.i) with ZIKV to determine if we could detect specific viral RNA species. Indeed, we observed full-length viral genomes that stained with all three probe sets (**Fig. 1B**, white arrow), partial degradation fragments that only stain with the 3’-CDS and sfRNA probes (**Fig 1B**, red arrow), or independent sfRNAs that lack staining with the 5’-CDS and 3’-CDS probes (**Fig. 1B**, cyan arrow). These data show that we can visualize specific viral RNA species, including sfRNAs.

We next analyzed the kinetics sfRNA production over the course of ZIKV infection by staining cells with the 3’-CDS and sfRNA probes. We observed that early during infection (4-8 hours p.i.), most sfRNA smRNA-FISH puncta co-stained with the 3’-CDS probes (**Fig. 1C**), indicating that they are either full-length genomes or longer 3’ degradation fragments. However, a small percentage of sfRNA smRNA-FISH puncta did not co-stain with the 3’-CDS, indicating that these are sfRNA molecules. By 12 hours p.i., total viral RNA increased and many sfRNA smFISH puncta lacked 3’-CDS staining, indicating the these are sfRNAs. The number of sfRNA smRNA-FISH puncta further increased by 18 hours p.i. and were diffusely localized throughout the cytosol (**Fig. 1C**).

Quantification of sfRNA and 3’-CDS smRNA-FISH puncta confirmed these observations by showing that the total number of cytosolic sfRNA puncta continually increased relative to the 3’-CDS puncta during infection (**Fig. 1C**). By 18 hours p.i., we detected ∼870 sfRNA smRNA-FISH puncta and ∼500 3’-CDS localized in the cytosol (**Fig. 1D**). We observed similar results during WNV infection, although quantification of individual sfRNAs was difficult due to the high abundance of WNV RNA (**Fig. S2**). These data demonstrate that sfRNA are produced early during flavivirus infection (6-18 hours p.i.) and diffusely localize throughout the cytosol.

### sfRNAs accumulate in cytoplasmic structures over the course of infection

An important observation we made is that by 24-48 hours p.i, 60-90% of A549 cells infected with DENV2, WNV, or ZIKV contained punctate cytoplasmic structures that stained for sfRNAs but not the 5’-CDS of the viral genomes (**Fig. 2A,B** and **Fig. S1A, S2A**). Co-staining ZIKV-infected cells with all three probe sets revealed that the cytoplasmic puncta mostly enriched for sfRNAs (**Fig. 2C**, white arrow). While the puncta sometimes co-stained with 3’-CDS probes (**Fig. 2C**, cyan arrow), they did not enrich for 5’-CDS, indicating that the puncta lack full-length genomes (**Fig. 2C**). We obtained similar results during DENV2 and WNV infection (**Fig. S3B**). The high intensity and the larger size of these structures argue that these are not individual sfRNA molecules, but instead that these are multiple sfRNA molecules concentrating at a structure. We refer to these as sfRNA granules in the following sections.

**Fig. 2.**
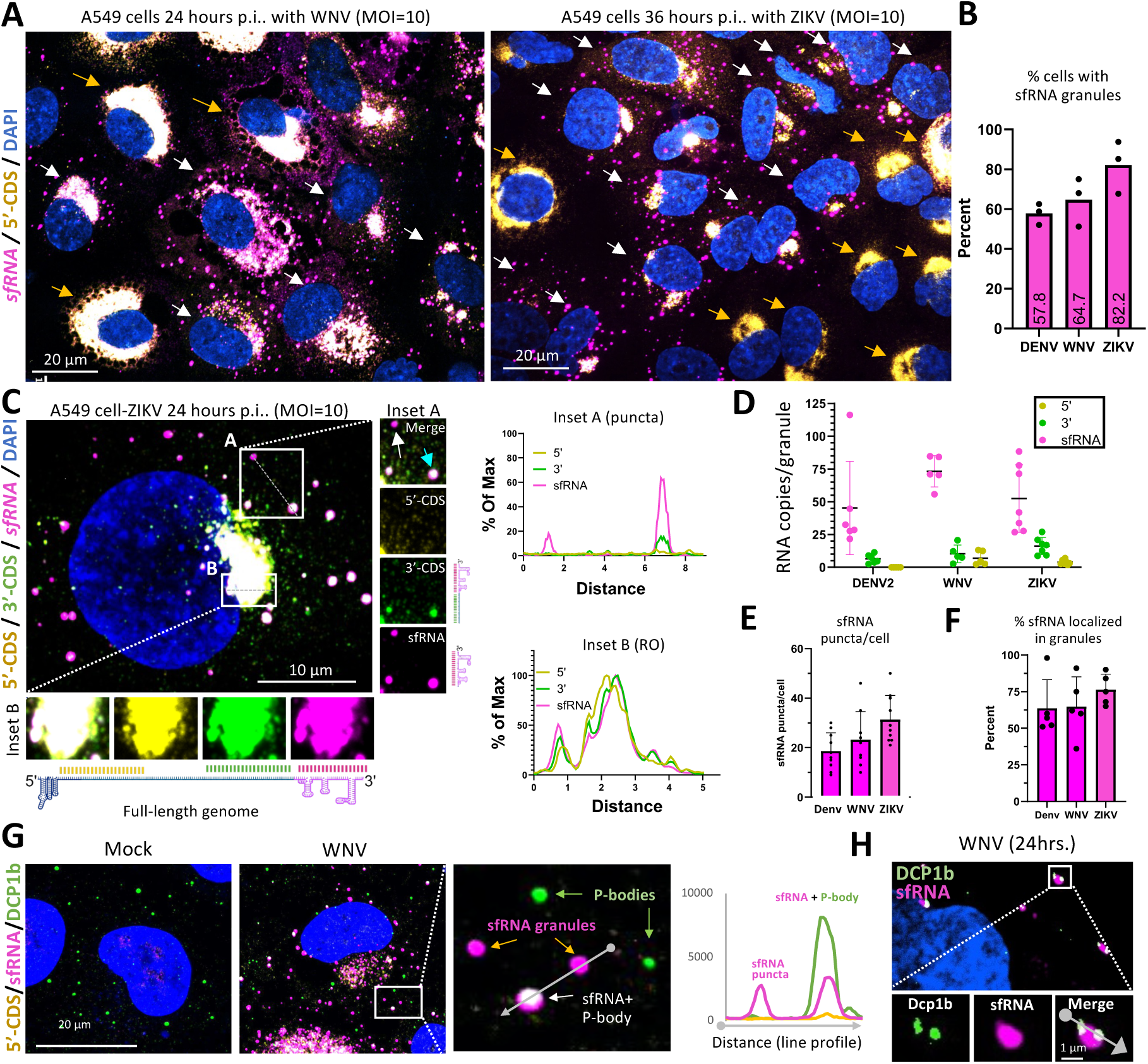
sfRNAs re-localize to cytoplasmic granules distinct from P-bodies. **(A)** Representative image of infected cells co-stained for 5’-CDS and sfRNAs 24 hours p.i. with WNV and 36 hours p.i. with ZIKV. White arrows correspond to cells containing sfRNA granules, while yellow arrows correspond to cells lacking these granules. **(B)** Percent of infected cells with sfRNA granules 24 hours p.i. in all three viruses. **(C)** Left: Co-staining smFISH for ZIKV 5’-CDS, 3’-CDS, and sfRNAs 24 hours p.i. of A549 cells with ZIKV (MOI=10). Right: Plotted profiles of lines in the insets **(D)** Estimated RNA molecules of each stain per sfRNA granule 24 hours p.i. **(E)** Number sfRNA granules per cell 24 hours p.i. **(F)** Percentage of total sfRNA (excluding the replication organelle) localized in granules at 24 hours p.i. **(G)** IF for DCP1b and smFISH for WNV 5’-CDS and sfRNAs 24 hours p.i. of A549 cells with WNV (MOI=10). Right of the inset, plotted profile shows raw intensity of indicated channels. **(H)** Super resolution microscopy of P-bodies (DCP1b) and WNV sfRNA granules.

Based on the intensity of individual smRNA FISH puncta vs. the intensity of sfRNA granules during ZIKV or WNV infection (**Fig. S3C**), each sfRNA granule contained an average of 50-75 copies of sfRNAs, 5-15 copies of longer 3’ fragments, and less than 10 RNAs containing the 5’-CDS (full-length genomes or 5’-end fragments) (**Fig. 2D**). These data indicate that sfRNAs, as well as some viral 3’-end fragments of flavivirus genomes, concentrate in sfRNA granules. We typically observed between 20-40 sfRNA granules per cell (**Fig. S3D)**. We estimate that between 1000-2000 sfRNA molecules accumulate in sfRNA granules per cell by 24 hours p.i. (**Fig. S3E**), which accounts for 60%-90% of total sfRNA molecules in the cell (**Fig. S3F**). We observed sfRNA granules form in primary human pulmonary artery endothelial cells infected with ZIKV (**Fig. S3G**), demonstrating that sfRNA granules form in non-cancer human cells.

We hypothesized that sfRNA granules are processing bodies (P-bodies) based on the rationale that sfRNAs associate with mRNA decay machinery that enrich in P-bodies.^9,16,17,34,35,36^ To test this, we first performed immunofluorescence (IF) assays against DCP1b, a P-body marker,^37,38^ and co-stained for sfRNAs. Although we observed instances in which P-bodies and sfRNA granules co-localized (**Fig. 2G**, white arrows), several observations indicate that sfRNA granules are not P-bodies.

First, WNV-infected cells contained twofold more sfRNA granules (median: 30) than DCP1b-positive P-bodies (median:14) (**Fig. 2G and Fig. S4A,B**). Second, only a small percentage (<20%) of sfRNA granules co-localized with P-bodies (**Fig. 2G** and **Fig. S4C**). Thus, the vast majority (>80%) of sfRNA granules do not co-localize with P-bodies. Third, only 30% of P-bodies stained for sfRNAs (**Fig. 2G** green arrows, **S4C**). Similar results were obtained in U-2 OS cells that stably express RFP-Dcp1a in both DENV2 and WNV infection (**Fig. S4D**). Lastly, super-resolution microscopy revealed that the sfRNA granules were often docked with P-bodies as opposed to being localized within P-bodies, with single sfRNA granules sometimes docking with multiple P-bodies (**Fig. 2H**, inset). We observed similar results during ZIKV infection, whereby sfRNA granules were distinct from P-bodies but were docked to P-bodies (**Fig. S4E**).

Combined, these data indicate that during the late phase of infection (24-48 hours p.i.) sfRNAs concentrate in cytoplasmic structures that are distinct from P-bodies but can interact with P-bodies.

### sfRNAs localize to RNase L-induced bodies

We hypothesized that sfRNA granules are either stress granules (SGs) or RNase L-induced bodies (RLBs). This is based on the rationale that both SGs and RLBs are ribonucleoprotein complexes that can assemble in the cytoplasm in response to viral infection, and both SGs and RLBs can interact with P-bodies.^29,30^ Additionally, flaviviruses are known to activate both PKR and RNase L,^39,40^ which trigger the assembly of SGs or RLBs, respectively.^27^

SGs and RLBs concentrate similar RNA-binding proteins, such as G3BP1.^27^ However, their assembly is mutually exclusive. This is because the activation of RNase L, which is required for RLB assembly, inhibits the assembly of SGs by degrading mRNAs that localize to SGs.^27^ For example, G3BP1 localizes to RLBs in wild-type (WT) A549 cells following lipofection of poly(I:C) (an immunogenic viral dsRNA mimic) (**Fig. 3A**). However, in RNase L-KO cells, lipofection of poly(I:C) results in SG assembly (**Fig. 3A**). Notably, SGs are more irregular in their morphology and larger (5μm diameter) in comparison to RLBs, which are smaller (1μm diameter) and more spherical (**Fig. 3A**). Additional key differences between SGs and RLBs is that SGs require PKR and G3BP1 for their assembly whereas RLBs do not, and RLBs do not typically enrich many SG-associated RBPs, such as TIA-1.^29^

**Fig. 3.**
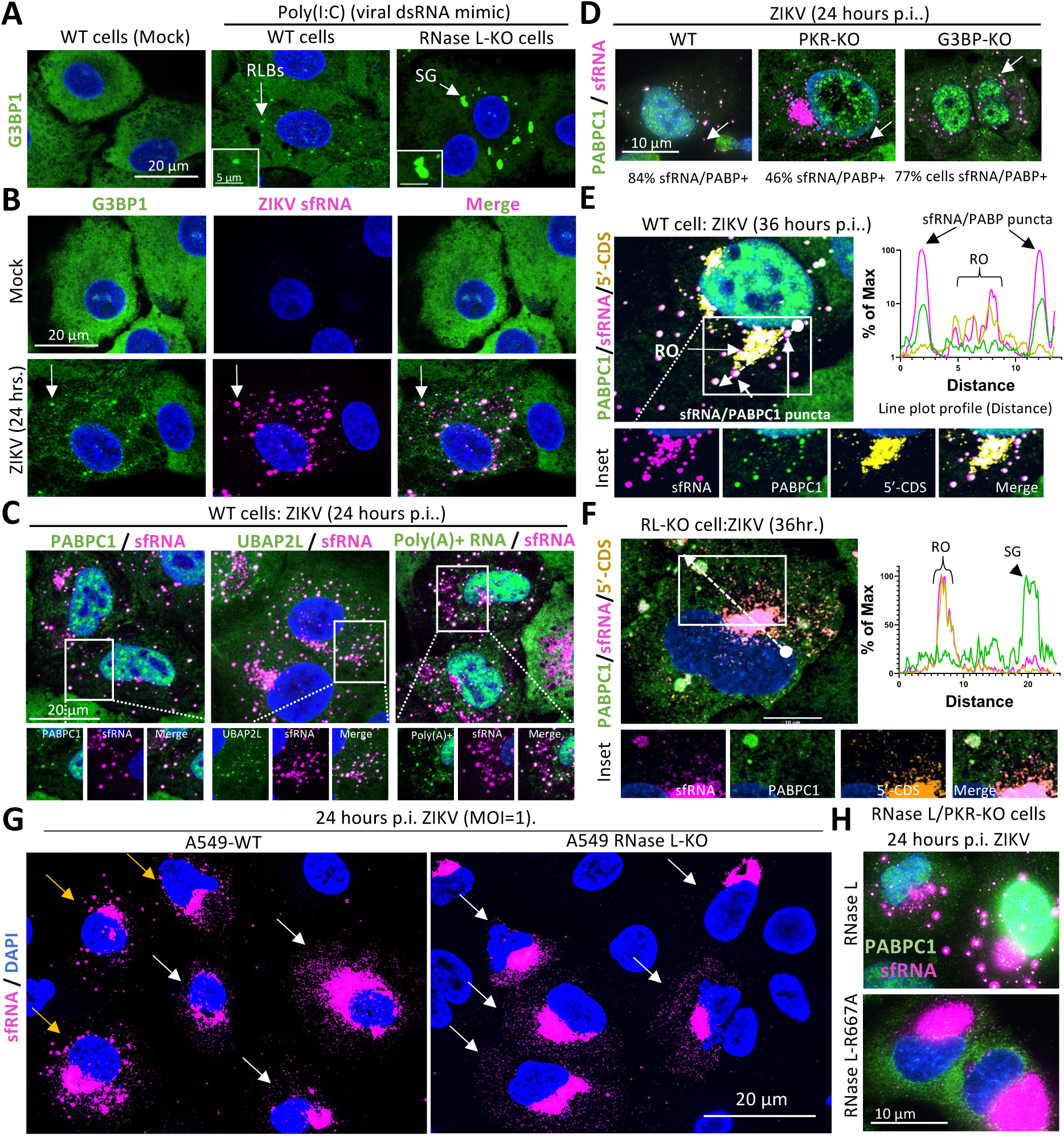
sfRNA localize to RNase L-induced bodies. **(A**) IF for G3BP1 in WT and RL-KO cells treated with Poly(I:C), demonstrating phenotypic differences between RLBs and SGs. **(B)** IF for G3BP1 and smFISH for ZIKV sfRNAs in A549 cells 24 hours p.i. (MOI=10) with ZIKV. (**C**) Similar to (**B**) but staining IF for PABPC1 or UBAP2L or RNA FISH for poly(A)+ RNA. Panels displaying individual stains are shown in **Fig. S5A-C**. (**D**) IF for PABPC1 and smFISH for ZIKV sfRNAs in parental (WT) A549 and RNase L, PKR and/or G3BP1/2 knockout A549 cells 24 hours p.i. with ZIKV (MOI=10). **(E-F)** PABPC1 IF and smFISH for ZIKV 5’ and sfRNAs in WT **(E)** and RLKO **(F)** A549 cells. The line graph on to the right of panels displays the intensity of PABPC1, sfRNA, and 5’-CDS staining from the line trace displayed in the merged image. **(G)** smFISH staining of ZIKV sfRNA in WT vs RLKO cells. Yellow arrows indicate cells with sfRNA puncta, white arrows indicate cells with no sfRNA puncta. **(H)** IF for PABPC1 and smFISH for ZIKV sfRNAs in PKR/RL-KO A549 cells rescued with either parental RL or a catalytically inactive version (R667A) 24 hours p.i. with ZIKV (MOI=10).

To determine if sfRNAs concentrate in either SGs or RLBs, we performed immunofluorescence assays for G3BP1 (RLB/SG marker) 24 hours p.i. with ZIKV. Importantly, we observed that nearly all ZIKV sfRNA granules co-stained for G3BP1 (**Fig. 3B**). Moreover, we observed additional RLB/SG markers – PABPC1, UBAP2L, poly(A)+ RNA – co-localize with sfRNA granules during ZIKV infection (**Fig. 3C** and **Fig. S5A-C**). PABPC1 and G3BP1 also co-localized with sfRNA granules during DENV2 and WNV infection (**Fig. S5D,E**). These data indicate that sfRNA granules are either RLBs or SGs.

Several observations indicate that the sfRNA granules are RLBs as opposed to SGs. First, the small and spherical morphology of sfRNA-G3BP1 granules in ZIKV-infected cells more closely resembled RLBs formed in WT cells as opposed to SGs assembled in RNase L-KO cells in response to poly(I:C) (**Fig. 3A,B**). Second, TIA-1 did not strongly enrich in sfRNA granules, thus suggesting these are RLBs as opposed to SGs (**Fig. S5F**). Third, sfRNA granules containing PABPC1 formed in PKR-KO (46% of cells) and G3BP1/2-KO (77% of cells) cell lines infected with ZIKV (**Fig. 3D**), which is a similar percentage observed in WT cells (**Fig. 2A**). In contrast, SGs that assemble in RNase L-KO cells in response to ZIKV infection were completely abolished upon knockout of PKR or G3BP1/2 (**Fig S5G**). Fourth, in WT A549 cells infected with ZIKV, sfRNA staining intensity in the PABPC1 puncta (RLBs), which lacked 5’-CDS staining, was 10-fold higher than the signal in the viral replication organelle (RO) (**Fig. 3E**). In contrast, neither ZIKV nor WNV sfRNAs highly enriched in SGs assembled in RL-KO cells (**Fig. 3F and S6A,B**).

Lastly, a key observation strongly supporting that sfRNA granules are RLBs is that they are dependent on RNase L for their formation since they do not form in RNase L-KO A549 cells infected with ZIKV, WNV, or DENV2 (**Fig. 3G, S6A-B**). Unlike in WT cells, in which ∼50% of ZIKV infected cells contained sfRNA granules 24 hours p.i. (**Fig. 3G**, yellow arrows), no RNase L-KO cells contained sfRNA granule structures (**Fig. 3G**, white arrows). Instead, sfRNA probes strongly stained the viral replication organelle (RO) or diffusely stained the cytoplasm (**Fig. 3G**, white arrows). Similarly, we did not observe sfRNA granules in WNV-infected c6/36 mosquito cells (**Fig. S7A**), which do not encode for RNase L and thus would not be expected to assemble RLBs. Because sfRNA granule formation is RNase L-dependent, this strongly supports that they are RLBs.

Importantly, the formation of sfRNA-PABPC1 granules in RNase L-null A549 cells infected with ZIKV was rescued cells upon stable expression of RNase L but not RNase L-R667A (catalytically inactive mutant) (**Fig. 3I**). These data indicate that the formation of sfRNA-PABPC1 granules in mammalian cells requires expression of catalytic competent RNase L, which is the key factor required for RLB assembly.^27^

Combined, these data argue that sfRNAs encoded by DENV2, ZIKV, and WNV localize to RNase L-induced bodies (RLBs) during viral infection in mammalian cells.

### sfRNAs re-localize to RLBs upon activation of RNase L-mediated RNA decay

We next sought to understand how sfRNAs localize to RLBs. RNase L is a latent endoribonuclease that is activated upon detection of viral dsRNA by oligoadenylate synthetase (OAS) proteins.^41^ Following activation, RNase L rapidly degrades cellular mRNAs (i.e., *GAPDH* mRNA) to completion.^27,42^ As cellular mRNAs degrade, RLBs assemble.^27^ The observation that not all A549 cells display sfRNA granules (**Figs. 2A and 3G**) led to the hypothesis that these cells had not activated RNase L and thus did not contain RLBs, whereas the cells with sfRNA granules activated RNase L and thus contained RLBs. To test this, we stained WT and RNase L-KO (RL-KO) A549 cells infected with ZIKV for human *GAPDH* mRNA, which is rapidly degraded upon activation of RNase L.^27^

Consistent with the hypothesis that sfRNAs concentrate at RLBs that form upon activation of RNase L-mediated RNA decay, we only observed sfRNA granules (RLBs) in ZIKV-infected A549 cells that did not contain *GAPDH* mRNA (**Fig. 4A**, white arrows, and **Fig. S7B**). In contrast, sfRNAs in ZIKV-infected A549-WT cells containing abundant *GAPDH* mRNA were diffusely distributed in the cytosol and did not contain sfRNA granules (RLBs) (**Fig. 4C,D**, yellow arrows). The degradation of *GAPDH* mRNA in response to ZIKV infection was dependent on RNase L because *GAPDH* mRNA levels were not reduced in ZIKV-infected RNase L-KO A549 cells (**Fig. 4A,B, S7B**). We obtained similar results during DENV2 and WNV infection (**Fig. S7C,D**). These data indicate that sfRNAs concentrate at RLBs following the activation of RNase L-mediated RNA decay and subsequent assembly of RLBs.

**Fig. 4.**
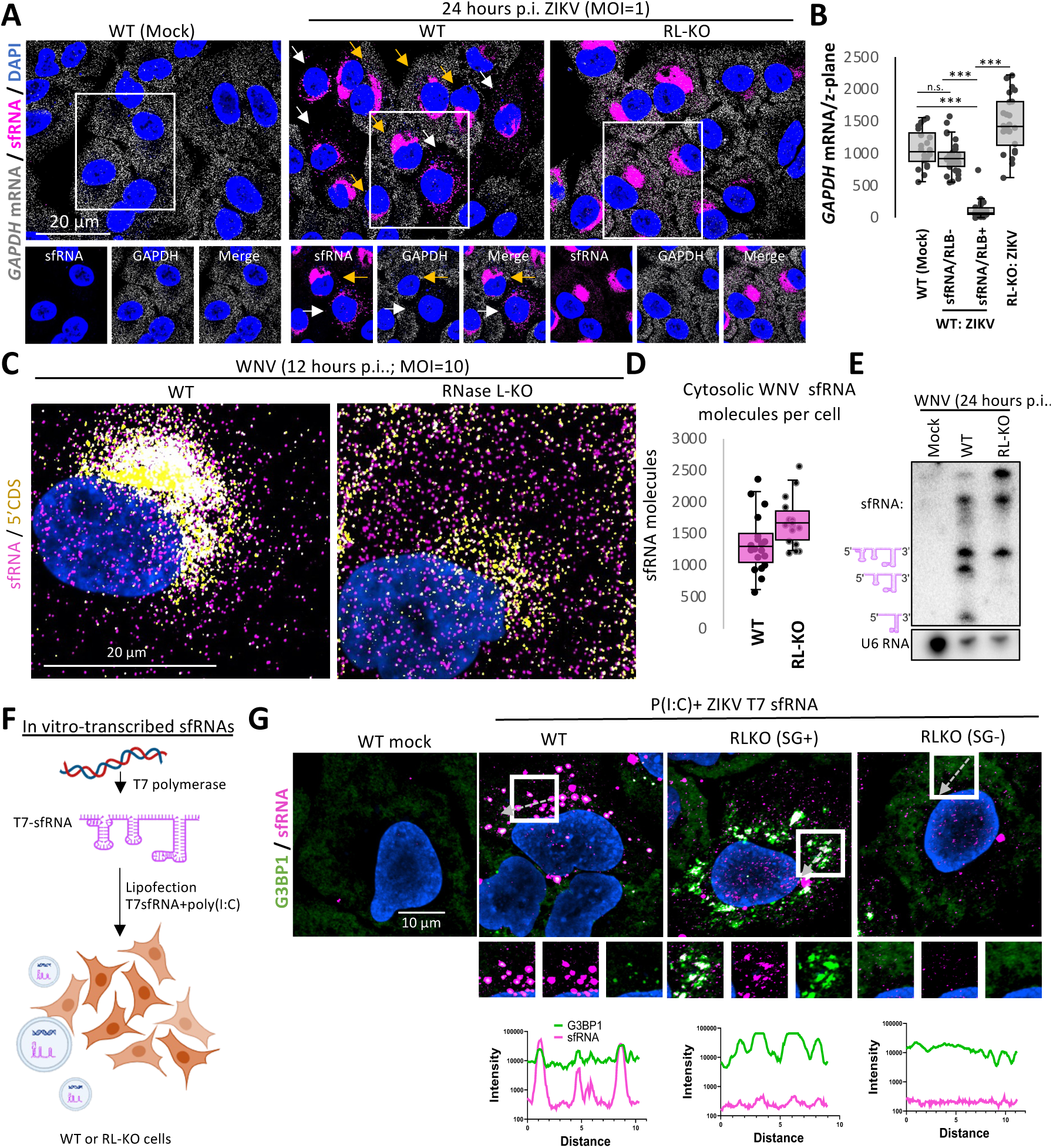
sfRNAs localize to RLBs following the initiation of RNase L-mediated mRNA decay. (**A**) smFISH for human *GAPDH* mRNA and ZIKV sfRNAs in mock or 24 hours p.i. with ZIKV (MOI=10) in parental (WT) and RNase L-KO (RL-KO) A549 cells. White arrows indicate cells with active RNase L, while yellow point to cells without RNase L activated. (**B**) Quantification of *GAPDH* mRNA smFISH in mock-infected WT cells, WT cells infected with ZIKV that did or did not display sfRNA granules, and RL-KO cells, as represented in (A). (**C**) smFISH for WNV 5’-CDS (yellow) and sfRNA (magenta) 12 hours p.i. (MOI=10). Individual magenta foci represent sfRNAs localized to the cytoplasm, whereas white dots (co-localized magenta and yellow foci) represent full-length genomes containing the 5’-CDS and sfRNA. The viral replication organelle (RO) is indicated, which concentrates full-length genomes. **(D)** Quantitation of sfRNA count by smFISH in WT vs RL-KO A549 cells at 12 hours p.i **(E)** Northern blot analysis for WNV sfRNAs 24 hours p.i. with WNV (MOI=10). **(F)** Schematic for generating sfRNA *in vitro* and transfecting into cells. **(G)** IF for G3BP1 and smFISH for ZIKV sfRNAs in co-transfected with or without in vitro-transcribed (IVT)-sfRNA (ZIKV) with poly(I:C) (RNase L activator) in WT and RNase L-KO cells with and without SGs. Gray arrows represent line used for line trace (below).

We considered two non-mutually exclusive possibilities by which sfRNAs localize to RLBs. First, sfRNAs that are generated by RNase L-mediated cleavage of viral RNA genomes shuttle to RLBs. In this scenario, RNase L would be expected to substantially increase overall sfRNA production considering 60-90% of sfRNA localize to RLBs. Second, sfRNAs are generated independently of RNase L and localize in the cytosol but, upon activation of RNase L, sfRNAs re-localize to RLBs. Several observations support that pre-formed sfRNA re-localize from the cytosol to RLBs, but that RNase L can promote and/or alter sfRNA production in a manner that could lead to incorporation of sfRNAs into RLBs.

RNase L has been proposed to specifically cleave viral RNA during the early phase of infection.^43^ Therefore, we considered the possibility that RNase L could specifically cleave viral RNAs to generate sfRNAs which then localize to RLBs. One prediction of this would be that sfRNAs levels would be lower in RNase L-KO cells. However, smRNA-FISH for WNV sfRNAs revealed a comparable number of sfRNA molecules are generated in WT and RNase L-KO A549 cells 12 hours p.i. (**Fig. 4C,D**). Thus, RNase L does not promote sfRNA generation by specifically cleaving viral genomes early during infection prior to widespread RNase L activation.

We next examined if widespread activation of RNase L, which occurs by 24 hours p.i (**Fig. S7B)**, increases WNV sfRNA production. Because viral RNAs are too abundant to reliably differentiate independent sfRNAs from full-length genomes at this time via smFISH (**Fig. S2A**), we performed northern blot analyses for WNV sfRNA. WNV sfRNA-1 (the largest sfRNA species) was comparable between WT and RL-KO cells (**Fig. 4E**), indicating that sfRNA-1 can be generated independently of RNase L. However, RNase L increased production of the smaller WNV-encoded sfRNA species (sfRNA-2 and sfRNA-3), presumably by cleaving in between dumbbell structures of sfRNA-1(**Fig. 4E**). Quantification of total sfRNA species indicates that RNase L increases sfRNA levels in cells by ∼twofold (**Fig**. **S8A**). Similar results were observed in ZIKV infection (**Fig. S8B)**. These data indicate that RNase L while RNase L increases steady state sfRNA levels, the small increase does not account for 90% of sfRNAs localizing to RLBs.

The observation that sfRNAs are abundant and diffusely localized throughout the cytosol in WT cells infected with ZIKV 24 hours p.i. that had not activated RNase L-mediated decay of cellular mRNAs indicates that sfRNAs are generated in high abundance prior to RNase L activation and then re-localize to RLBs upon RLB assembly. To test if sfRNAs could localize to RLBs independently of degradation of the full-length viral genome, we generated ZIKV sfRNA-1 using T7 polymerase (T7-sfRNAs). Following purification, we co-transfected ZIKV T7-sfRNA-1 into WT or RNase L-KO A549 cells with poly(I:C) to activate RNase L (**Fig. 4F**). In WT cells that activated RNase L and assembled RLBs, sfRNAs were mostly localized to RLBs, whereas in RNase L-KO cells, which do not form RLBs, the T7-sfRNAs were widely distributed in the cytosol (**Fig 4G**). We note that T7-sfRNAs also localized to the SGs in RL-KO cells that generated SGs, though the enrichment of T7-sfRNA in SGs was ∼10-fold less than those observed in RLBs (**Fig. 4G**).

Taken together, these data indicate that pre-formed sfRNAs re-localize from the cytosol to RLBs following activation of RNase L-mediated RNA decay and subsequent RLB assembly. However, we cannot rule out that the cleavage of sfRNA-1 by RNase L results in sfRNA-2 and sfRNA-3 incorporation into RLBs.

### RLBs sequester sfRNAs away from mRNA decay machinery and P-bodies to promote viral RNA decay

We next wanted to understand how RLB-mediated sequestration of sfRNAs alters the cellular response to flavivirus infection. A primary function of sfRNAs is the inhibition of cellular mRNA decay. This function is supported by studies showing that sfRNAs interact with mRNA decay machinery^17,44^ and can localize to P-bodies, which enrich for mRNA decay machinery.^9^ Moreover, sfRNAs can limit XRN1-mediated decay of cellular and viral RNAs. Thus, we hypothesized that RLBs could sequester sfRNAs away from cellular RNA decay machinery in the cytoplasm and/or P-bodies, and this in turn would promote decay of cellular and viral RNA. The following observations support this hypothesis.

#### RLB assembly shunts WNV sfRNAs from P-bodies to RLBs

We considered the possibility that sfRNA sequestration by RLBs could reduce sfRNA association with mRNA decay machinery. One prediction of this hypothesis is that in the absence of RLBs, sfRNAs would enrich in P-bodies. However, upon RNase L activation and RLB assembly, sfRNA association with P-bodies would decrease. To test if RLBs could reduce sfRNA interactions with mRNA decay machinery in P-bodies, we stained infected WT and RNase L-KO cells with WNV and stained G3BP1 (RLB marker), DCP1b (P-body marker), and sfRNAs. In RNase L-KO cells, which do not assemble RLBs based on the lack of G3BP1 puncta, we observed that WNV sfRNAs strongly enriched in all P-bodies by 24 hours p.i. (**Fig. 5A,B**). These data indicate that WNV sfRNAs associate with mRNA decay machinery that can enrich in P-bodies.

**Fig. 5.**
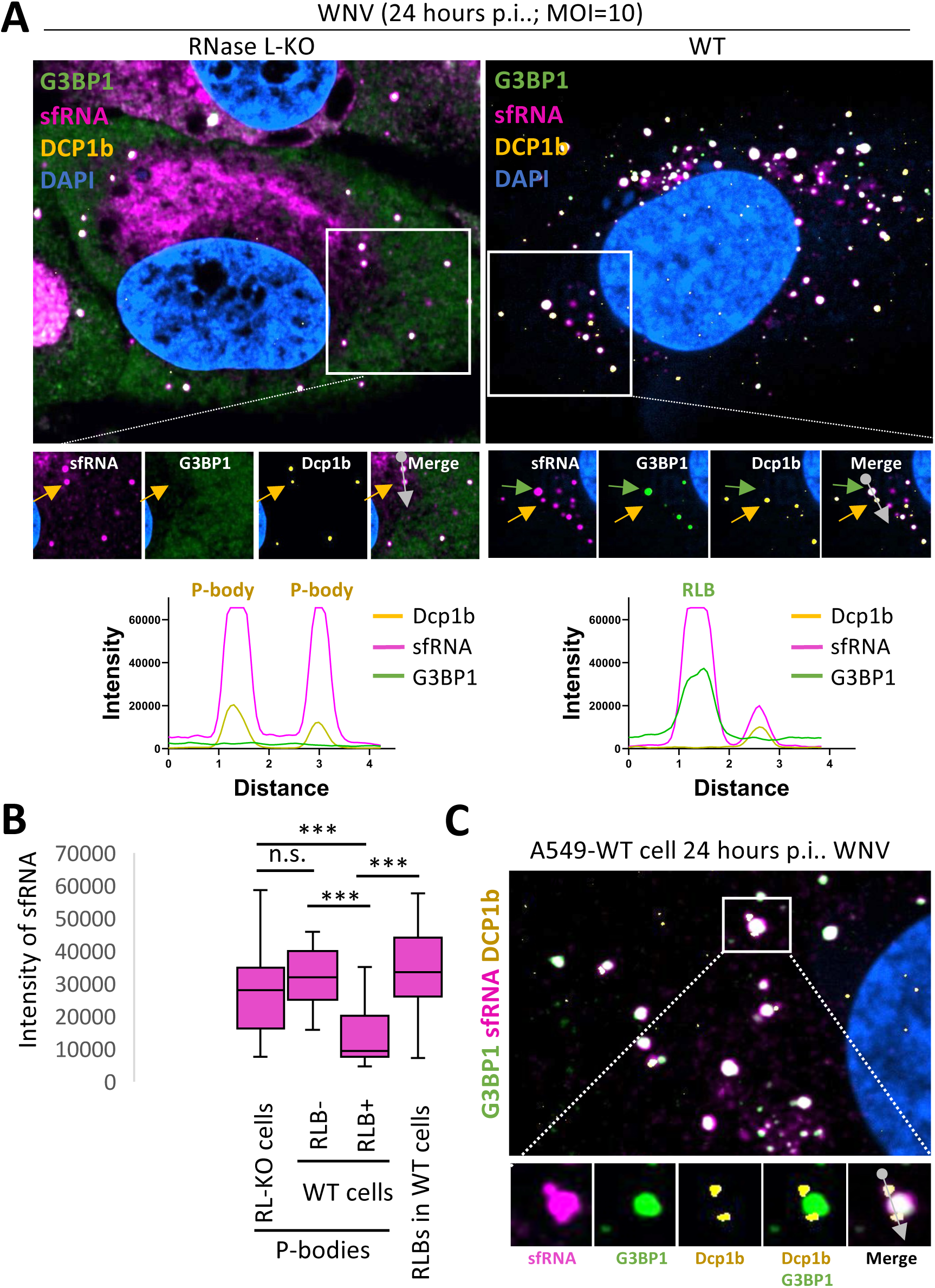
sfRNAs re-localize from P-bodies to RLBs upon RNase L activation. **(A)** IF of G3BP1, Dcp1b and smFISH of sfRNA in RLKO and WT A549 cells. Grey lines indicate plot profile (below). **(B)** Mean intensity of sfRNAs in P bodies in binned cell types. **(C)** Super-resolution of IF for G3BP1 and Dcp1b co-stained with sfRNA. Inset and line trace show example of docked P body.

Importantly, in WT A549 cells that assembled RLBs, sfRNA intensity in P-bodies was ∼3.5-fold lower than those observed in P-bodies in RNase L-KO cells (**Fig. 5A,B**). Moreover, sfRNA intensity was 3.5-fold higher in RLBs than P-bodies in these WT cells (**Fig. 5A,B**). Quantification of sfRNA enrichment in P-bodies in RNase L-KO cells or in WT cells that either contained (RLB+) or did not contain RLBs (RLB-) in several cells confirmed these results. In comparison to WT cells that lacked RLBs or RNase L-KO cells, sfRNA enrichment in P-bodies was significantly reduced in WT cells that contained RLBs (**Fig, 5B**). Importantly, we observed a corresponding increase in WNV sfRNAs enriched in RLBs (**Fig. 5B**), indicating that sfRNAs shunt from P-bodies to RLBs. Notably, we observed docking between sfRNA-positive P-bodies and RLBs (**Fig. 5C**), suggesting that sfRNAs could be directly transferred from P-bodies to RLBs.

#### XRN1-mediated decay of cellular mRNAs is robust in flavivirus-infected cells containing RLBs

sfRNAs are known to inhibit XRN1-mediated decay of cellular mRNAs.^17,45^ However, two observations suggest that flavivirus-infected cells that contain RLBs have robust cellular mRNA decay potential. First, smFISH analyses showed robust decay of *GAPDH* mRNA upon activation of RNase L during DENV2, ZIKV, or WNV infection (**Fig. 4A and Fig. S7**). Consistent with these data, total poly(A)+ RNA was significantly reduced in an RNase L-dependent manner in ZIKV- and WNV-infected cells containing RLBs (**Fig. 6A,B and Fig S9A-B**). These data demonstrate that upon RNase L activation, flavivirus-infected cells have robust cellular mRNA decay potential.

**Fig. 6.**
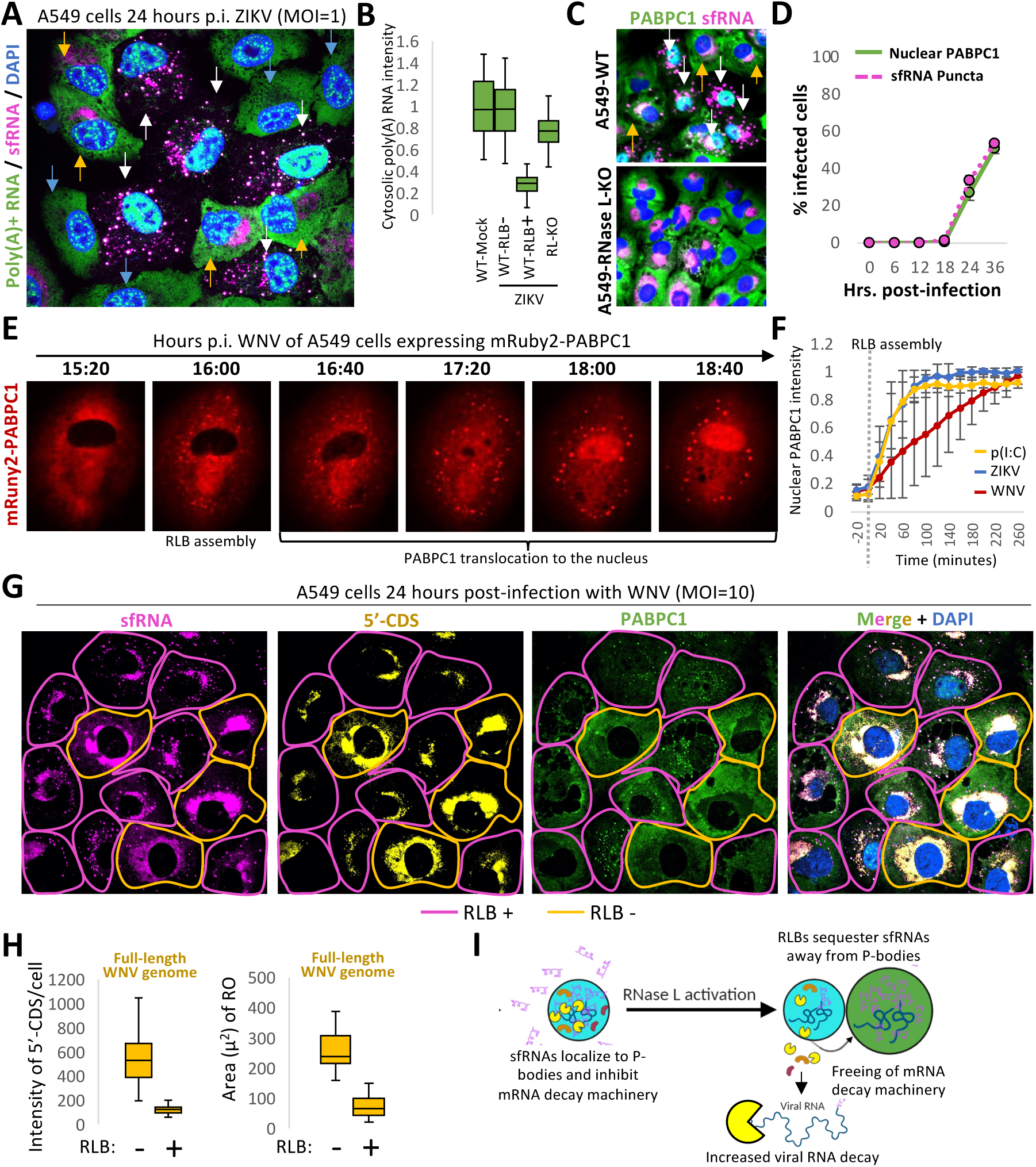
RLBs sequester sfRNAs to re-establish XRN1-mediated decay of viral RNA. (**A**) FISH for Poly(A) RNA and smFISH for ZIKV sfRNAs at 24 hours p.i. (**B**) Quantification of the cytosolic levels of Poly(A) RNA in listed cell types during ZIKV infection. (**C**) IF of PABPC1 and smFISH of sfRNA in WT and RL-KO A549 cells. White arrows point to cells with PABC1 translocation into the nucleus, yellow point to cells lacking nuclear PABC1. (**D**) Quantification of images from (C) **(E)** Live-cell imaging of A549 cell expressing mRuby2-PABPC1 infected with WNV virus. **(F)** Single-cell quantification of PABP translocation rate during poly(I:C), ZIKV and WNV. Zero timepoint set to first RLB formation in each cell. **(G)** Co-staining of smFISH of sfRNA, 5’-CDS, and IF for PABPC1 24 hours p.i. Pink outlines indicate RNase L activation, while yellow outlines indicate inactive RNase L. **(H)** Quantification of intensity and size of replication organelle in cells with or without RNase L active from (G). **(I)** Proposed model of sfRNA function.

Second, when RNase L cleaves cellular mRNAs and XRN1 degrades the 3’-fragment, poly(A) binding protein C1 (PABPC1) is released from poly(A) tails and translocates to the nucleus.^27^ Importantly, we observed RNase L-dependent PABPC1 translocation to the nucleus in WNV- and ZIKV-infected cells that contained RLBs (**Fig. 6C and Fig. S10A**), and that the initiation of PABPC1 translocation to the nucleus over the course of infection closely coincided with sfRNA-RLB formation (**Fig. 6D** and **Fig. S10B,C)**

We then performed live-cell imaging of PABPC1 translocation during WNV or ZIKV infection or following poly(I:C) lipofection to determine the rate PABPC1 translocation (**Fig. 6E and Fig. S11A,B**). These data demonstrated that the rate of PABPC1 translocation following RLB formation between ZIKV-infected cells and poly(I:C)-lipofected cells was comparable (**Fig. 6F and Fig. S11A-C**). WNV-infected cells displayed a slightly slower rate of PABPC1 translocation compared to ZIKV and poly(I:C) (**Fig. 6E and Fig. S11C)**, though it did fully translocate within 4 hours following RLB assembly. Notably, WNV sfRNAs more highly enrich in P-bodies prior to RNase L activation in comparison to ZIKV sfRNAs, which do not highly enrich in P-bodies in cells with inactive RNase L (**Fig. S11F**). We interpret these data to indicate that WNV sfRNAs more strongly antagonize mRNA decay machinery, thus resulting in a reduction in mRNA decay kinetics until RLBs can sequester WNV sfRNAs from P-bodies.

We confirmed that PABPC1 translocation to the nucleus during viral infection is XRN1-dependent by showing that knockdown of XRN1 reduced the rate of PABPC1 translocation during ZIKV infection (**Fig. S11A**). These data demonstrate XRN1 is active in cells that have activated RNase L-mediated assembly of RLBs.

#### RLB assembly correlates with reduced viral genomic RNA

The observation that XRN1 is active in cells that have activated RNase L and formed RLBs led us to ask whether RLBs could promote viral RNA decay. To test this, we measured 5’-CDS staining (full-length genome) in A549 cells infected with WNV 24 hours p.i. We stained for PABPC1 and sfRNAs to identify cells that activated RNase L and thus contained sfRNA+ RLBs and nuclear PABPC1. We then quantified 5’-CDS in cells that contained RLBs (RLB+) or did not contain RLBs (RLB-).

Importantly, we observed a significant reduction (∼5-fold) in full-length genome (5’-CDS staining) in WNV-infected WT A549 cells that contained RLBs in comparison to cells that lacked RLBs (**Fig. 6G,H**). Moreover, we observed a significant reduction (3-fold) in the area of the replication organelle (RO) (**Fig. 6,G,H**). Similar results were obtained with ZIKV (**Fig. S12A,B**). These data further support that RLB-mediated sequestration of sfRNA can promote the decay of viral mRNAs/genomes by cellular RNA decay machinery (**Fig. 6I**).

## DISCUSSION

In this article, we analyzed the production and localization of sub-genomic flavivirus RNAs (sfRNA) by smRNA-FISH during DENV2, WNV, or ZIKV infection (**Fig. 1**). We observed that the vast majority (∼90%) of sfRNAs generated by DENV2, WNV, and ZIKV re-localize from the cytosol to cytoplasmic ribonucleoprotein complexes termed RNase L-induced bodies (RLBs) (**Figs. 1,2**), which form in infected cells when the RNase L-mediated RNA decay pathway is activated (**Figs. 3-4**). In cells that assemble RLBs, we observed reduced localization of WNV sfRNAs in P-bodies (**Figs. 5**), and this coincided with an increase in decay of viral RNA via XRN1 (**Fig. 6**). Based on these observations and previous literature supporting that sfRNAs inhibit cellular RNA decay,^17,44,45^ we propose that the RLBs sequester sfRNAs away from RNA decay machinery localized in the cytosol and P-bodies, thus leading to an increase in the capacity for cellular RNA decay pathways to degrade viral RNA (**Fig. 6I**).

The identification of subgenomic flavivirus RNAs localizing to RLBs is important because specific cellular RNAs had yet to be detected in RLBs, and viral RNAs were not known to localize to RLBs. While previous studies failed to identify full-length cellular mRNAs in RLBs,^27,29,30^ RLBs strongly enrich for cellular poly(A)+ RNA.^27,29^ This has led to the hypothesis that RLBs concentrate the degradation fragments of mRNAs containing poly(A)+ tails following the activation of RNase L-mediated mRNA decay.^30^ Thus, RLBs concentrating subgenomic viral RNAs (sfRNAs and longer 3’-end fragments) but not full-length viral genomes supports the notion that RLBs are composed of 3’-end fragments of RNAs generated via cellular RNA degradation pathways following RNase L activation. Based on these findings, RLBs appear to be involved in the turnover of host/viral mRNAs and/or are sites to compartmentalize host/viral mRNAs that cannot be fully degraded by typical RNA decay pathways.

Our data indicate that two non-mutually exclusive mechanisms potentially contribute to sfRNA localization to RLBs. The first potential mechanism is that RNA fragments, such as sfRNAs, are generated by RNase L-mediated cleavage of longer RNA, such as viral genomes, specifically shuttle to RLBs. While RNase L is not required for sfRNA-1 generation based on our northern blot analyses 24 hours p.i. with either WNV or ZIKV (**Fig. 4E** and **Fig. S8**), RNase L was required for the biogenesis of downstream sfRNA fragments, sfRNA-2 and sfRNA-3 (**Fig 4E** and **Fig. S8B**). Thus, it is possible that the cleavage of sfRNA-1 into sfRNA-2 and sfRNA-3 by RNase L results in their incorporation into RLBs.

A second potential mechanism by which sfRNA localize to RLBs is through sfRNA interactions with RNA-binding proteins that concentrate in RLBs, which would result in sfRNA re-localization from the cytosol to RLBs upon RLB assembly. Consistent with this, we observed sfRNAs widely distributed in the cytosol early during infection prior to RNase L activation (**Fig. 1C**) in cells with inactivate RNase L (**Fig. 4A**). However, in cells with RLBs, nearly all sfRNAs localize to RLBs, suggesting that sfRNAs re-localize to RLBs after RLB assembly. Consistent with this, pre-formed T7-generated sfRNAs localized to RLBs (**Fig. 4G**). This suggests that sfRNAs interact with an RNA-binding protein that concentrates in RLBs. Notably, we observed a low level of sfRNAs enrich in SGs that formed in RNase L-KO cells in response to viral infection or T7-sfRNA lipofection (**Fig. 3F** and **Fig. 4G**). While sfRNAs did not greatly enrich in SGs, which is consistent with the fact that short RNAs do not typically concentrate in SGs ^46–48^, this suggests that an RNA-binding protein common to RLBs and SGs may be responsible for sfRNA localization to RLBs. Although sfRNAs have been shown to bind G3BP1^49^, G3BP1 is not the RNA-binding protein responsible for this based on the observation that sfRNAs localize to RLBs in G3BP1/2- KO cells (**Fig. 3D**). Studies are underway to further define the molecular basis responsible for sfRNA sequestration in RLBs.

Previous studies have shown that sfRNAs interact with cellular RNA decay factors (XRN1, EDC3, DDX6) that enrich in P-bodies^9,34,45,50^. Moreover, FISH analyses of sfRNAs showed that sfRNAs primarily co-localize with XRN1 in P-bodies in A549 cells^9^. Thus, a consensus model for the function of sfRNA is that they antagonize cellular RNA decay pathways to increase the stability of viral RNA by inhibiting mRNA decay machinery that enrich in P-bodies. Our data is consistent with these previous findings, as we observed sfRNAs localize to P-bodies in cells with inactive RNase L **(Fig. 5A**). However, our data adds to these studies by showing that most sfRNAs re-localize to RLBs in cells that activated RNase L and assembled RLBs, with RLBs sequestering most of the sfRNA that would otherwise associate with mRNA decay machinery in P-bodies (**Fig. 5**). The interactions between RLBs and P-bodies suggest that sfRNAs might be directly transferred from P-bodies to RLBs (**Fig. 5C**), although re-partitioning of cytosolic sfRNAs to RLBs could also lead to a reduction of sfRNAs in P-bodies over time (**Fig. 5A,B**). Nevertheless, the reduction of sfRNA localization to P-bodies in cells with RLBs suggests that sfRNAs are no longer associating with RNA decay machinery, which would be expected to promote RNA decay machinery.

Three observations indicate that RNA decay machinery is active in mammalian cells that contain RLBs despite the presence of sfRNAs. First, *GAPDH* mRNAs and total poly(A)+ RNA decreased in cells that have activated RNase L and assembled RLBs (**Fig. 4A** and **Fig. 6A**). Second, PABPC1 rapidly translocates from the cytosol to the nucleus in an XRN1-dependent manner following the activation of RNase L and RLB formation in cells infected with DENV2, ZIKV, or WNV (**Fig. 6C** and **Fig. S11D**).^51^ Lastly, we observed enhanced decay of viral genomes/mRNAs in ZIKV- and WNV-infected cells that contained RLBs (**Fig. 6G,H** and **S12**), indicating that cellular RNA decay machinery is functional in cells in which sfRNAs have been sequestered. A limitation of these data is that RNase L can cleave viral RNAs and thus could account for this reduction in overall viral RNA levels. However, we would expect that RNase L-mediated cleavage would not result in a decrease in total fluorescence of viral RNAs without further decay by XRN1. Thus, we interpret our data to suggest that XRN1 is functional in degrading viral genomes following RLB assembly, and that both RNase L-mediated cleavage of viral genomes and RLB-mediated sequestration of sfRNA promotes XRN1-mediated decay of viral genomes. Current studies are underway to identify factors other than RNase L that are required for RLB assembly, which will allow for further dissection of the function of RLBs.

Taken together, our data strongly argue that the OAS/RNase L antiviral pathway regulates sfRNA biology in mammalian cells, both by altering sfRNA biogenesis and by reducing the interactions of sfRNAs with RNA decay machinery via sequestration to RLBs. While our data is consistent with RLB-mediated sequestration of sfRNAs promoting decay of viral RNA, we do not rule out the possibility that sfRNAs could inhibit a yet-to-be identified function of RLBs, or that RLBs could inhibit additional functions of sfRNA. Ongoing studies aim to further understand the molecular basis and functional consequences of sfRNA and RLB interactions, as well as identify additional viral RNAs that localize to RLBs.

## ACKNOWLEDGMENTS

We thank Dr. Roy Parker (University of Colorado Boulder) for providing valuable commentary. We thank Dr. Hyeryun Choe and Dr. Lizhou Zhang (Boston Children’s Hospital) for providing viral stocks. This work was funded by institutional funds from The Herbert Wertheim University of Florida Scripps Institute for Biomedical Innovation and Technology (JMB) and by the National Institute of General Medical Sciences of the National Institutes of Health under Award Number R35GM151249. The content is solely the responsibility of the authors and does not necessarily represent the official views of the National Institutes of Health.

## AUTHOR CONTRIBUTIONS

Conceptualization: JMW, JMB

Methodology: JMW, JMB

Investigation: JMW, JMB

Visualization: JMW, JMB

Funding acquisition: JMB

Project administration: JMB

Supervision: JMB

Writing – original draft: JMW, JMB

Writing – review & editing: JMW, JMB

## DATA AND MATERIALS AVAILABILITY

All data, code, and materials used in the analysis will be made available upon request to JMB.

## MATERIALS AND METHODS

### Cell Culture

A549 and U2OS cells were cultured in DMEM (Corning, cat#10-013-CV) +10% HI FBS(Sigma Aldrich, cat#F4135-500ML) + penicillin/streptomycin (Sigma, cat #P4333). Passaging used 0.25% trypsin (Corning, cat#45001-082). C6/36 and Vero cell lines were cultured in EMEM (cat #30-2003) + 10% FBS + P/S. CCL125 cells were cultured in EMEM +20% FBS + P/S. C6/36 and CCL125 cells were cultured at 28C with 5% CO2. A549, Vero, and U2OS cells were cultured at 37C with 5% CO2.

### Generation and quantification of viral stock

Viral stocks were produced in Vero cells. Once the cells were >80% confluent, cells were infected at 0.1 MOI. Supernatants were harvested at either 48 or 72 hours. After collecting the supernatant, dead cells were pelleted by centrifugation at 500g for 5 minutes. The remaining media was syringe filtered using a 0.45 uM filter.

### Viral infections

Cells were plated one day before infection and grown to 60-80% confluency. Viruses were added in serum-free media at indicated MOIs. After one hour, media was removed, cells were washed 1x with Dulbecco’s PBS (VWR, cat#02-0119-1000), and fresh media was added.

### smFISH probe labeling

The smFISH probes were labeled with Atto-488, Atto-550, or Atto-633 using 5-Propargylamino-ddUTP-ATTO-488 (Axxora: JBS-NU-1619-488), 5-Propargylamino-ddUTP - ATTO-550 (Axxora: JBS-NU-1619-550), or 5-Propargylamino-ddUTP-Atto633 (Axxora: JBS-NU-1619-633) and terminal deoxynucleotidyl transferase (Thermo Fisher Scientific: EP0161) as described in Gaspar et al. (2017). Oligo d(T)30-Cy3 were purchased from IDT.

### Microscopy

Cells were plated onto glass coverslips (Fisher Scientific Co LLC:NC1418755) in 12-well plates(Greiner Bio-One:82050-930), then virally infected. After the indicated time of infection, the media was removed, cells were washed 1x in DPBS, and then fixed in 500uL 4% paraformaldehyde for 12 minutes. To permeabilize the cells, pfa was then removed, and 1mL 75% ethanol was added. Cells were then stored at 4C for at least two hours before beginning staining protocol.

For dual staining of immunofluorescence and smFISH, cells were washed 2x in PBS, then incubated in 500uL primary antibody in PBS for four hours at 4C. Cells were washed 2x in PBS, then secondary antibodies were added for two hours at 4C. Cells were washed 2x, then fixed in 4% pfa (Fisher Scientific Co LLC: 50980495) for 10 minutes. Cells were washed 3x in PBS, then washed in buffer A (filter-sterilized 2x SSC with 10% formamide) for 5 minutes. smFISH probes were then added to a hybridization chamber (square petri dish cat #) containing a wet paper towel with parafilm on top. 50uL of smFISH probes diluted 1:100 in hybridization buffer (0.45um filtered(Fisher Scientific Co LLC: 09-719D), 10% dextran sulfate(Fisher Scientific Co LLC: S4030), 10% formamide (Fisher Scientific Co LLC: BP227500), 1x nuclease-free SSC(Life Technologies Corporation: 15557044), diluted in nuclease-free water(Fisher Scientific Co LLC: 10977023)) were dropped onto the parafilm. Glass slips were then flipped onto the smFISH probes. Hybridization chambers were sealed with parafilm and incubated overnight at 37C. The next day, slips were washed 2x in Buffer A, then once in 2x SSC. Slips were then mounted on slides (cat #) with Vectashield ((Vector Laboratories: 101098-044) and dried. Images were taken on a Nikon Eclipse Ti2 with a CFI60 Plan Apochromat Lambda D 100x Oil Immersion Objective Lens, N.A. 1.45, W.D. 0.13mm, F.O.V. 25mm, DIC, Spring Loaded. The filter set included: C-FL DAPI Filter Set, High-Signal-Noise, Semrock Brightline®, Excitation: 356/30nm (341-371nm),

### RNA extraction

After infection, 1mL Trizol (Life Technologies: 15596018) was used to resuspend each well of a 6 well plate. All remaining steps were at 4C or on ice. 200uL chloroform (Fisher Scientific: BPC298500Z) was added, then tube was mixed by inversion. Tubes were centrifuged for 15 minutes at 13,000 rcf. Aqueous phase was slowly removed and added into a new tube. 400uL of 2-propanol (Fisher Scientific: BPA451SK1) was added. Tubes were mixed, then spun for 15 minutes at 13,000 rcf. Liquid was aspirated from pellet, then pellet was washed in 500uL 75% ethanol (Fisher Scientific Co LLC: B09SBN6QQJ). Samples were centrifuged 13,000 rcf for 5 minutes, then ethanol was removed. Any remaining ethanol was evaporated off on a heat block at 65C. Pellets were then resuspended in 50uL nuclease-free water.

### Northern Blot protocol

2-10ug of RNA extractions were loaded into a 5% acrylamide (Bio-Rad Laboratories: 1610156), 0.5x TBE (Bio-Rad: 1610733) gel. Samples were run at 150V for 1 hour. Images were taken with ethidium bromide for visualization of RNAs. Gel was then transferred to BrightStar-Plus Positively Charged Nylon Membrane (Life Technologies: AM10102) at 30V for 90 minutes. Membrane was dried, then hybridized using a HL-2000 HYBRIDIZATION OVEN (Fisher: UVP95003101), and put in NorthernMax Prehybridization/Hybridization Buffer (Life Technologies: AM8677) for 1 hour. Probes were labeled using ATP[γ-32P]- 6000Ci/mmol 10mCi/ml EasyTide Lead, 250 µCi (PerkinElmer Health Sciences: NEG502Z250UC) and T4 PNK (NEB: M0201S) for 1 hour in a 37C water bath. Probes were added to blot in hybridization buffer overnight. Blots were then washed in 2x SCC, 0.1% SDS buffer 3x, then exposed to a phosphor screen (cat#) overnight. Imaging was performed on a XXX machine.

### T7 Transcription and transfection

T7 constructs were ordered from IDT and synthesized using HiScribe T7 High Yield RNA Synthesis Kit (NEB, #E2040S). RNA was purified via RNA extraction, as previously described, and quantitated via nanodrop. Tranfections then followed protocols for lipofectamine 2000 (Invitrogen: 11668019), using 1ug/well poly(I:C) and 500ng/well synthetic RNA in 12-well plates.

### Quantification of sfRNA granules

Quantification was performed using NIS-Elements AR. Percentage of cells containing sfRNA granules were manually counted. To measure the total percentage of sfRNA that colocalized to PABP granules, individual PABP puncta were circled as regions of interest, then the total fluorescence of those puncta was measured. Total sfRNA was measured by circling entire cells, then subtracting the fluorescence in the replication organelle, which was also manually circled.

### smFISH probes

GAPDH smFISH probe (Biosearch Technologies Inc.: SMF-2026-1). All other probes were generated using Stellaris probe design tool.

### FISH probes

Poly(A) RNA was detected using oligo(dT)18 Cy5, ordered from IDT.

Antibodies used (table)

*Antibodies used (table)*
| **Primary Antibodies** | **Vendor** | **Cat #** |
| --- | --- | --- |
| *DCP1B (D2P9W) Rabbit mAb* | *Cell Signaling Technology Inc* | 13233S |
| *XRN1 Polyclonal Antibody* | *Life Technologies Corporation* | A300443A |
| *Anti-PABP antibody* | *Abcam* | ab21060 |
| *Anti-UBAP2L antibody* | *Abcam* | ab70319 |
| *G3BP1 Polyclonal Antibody* | *Life Technologies Corporation* | PA529455 |
| **Secondary Antibodies** |  |  |
| Goat pAb to Rb IgG Alexafluor 647 | *Abcam* | Ab150079 |
| Goat Anti-Rabbit IgG H&L (FITC) | *Abcam* | AB6717 |
| Goat pAb to Ms IgG (Alexafluor 555) | *Abcam* | AB150114 |
| Goat Anti-Mouse IgG H&L (FITC) | *Abcam* | AB6785 |

### Figure generation

Models were created using BioRender.com. Line tracings and graphs were created using Graphpad Prism and Microsoft Excel.

### Statistical analyses

P-values were derived either by student’s t-test (Excel). The specific test used for each figure is specified in each figure legend. * p<0.05, **p>0.01, ***p>0.001 unless otherwise noted in the figure legend.

