## Supplemental Figures for "RNase L-induced bodies sequester subgenomic flavivirus RNAs and re-establish host RNA decay"

### SUPPLEMENTAL FIGURES AND LEGENDS

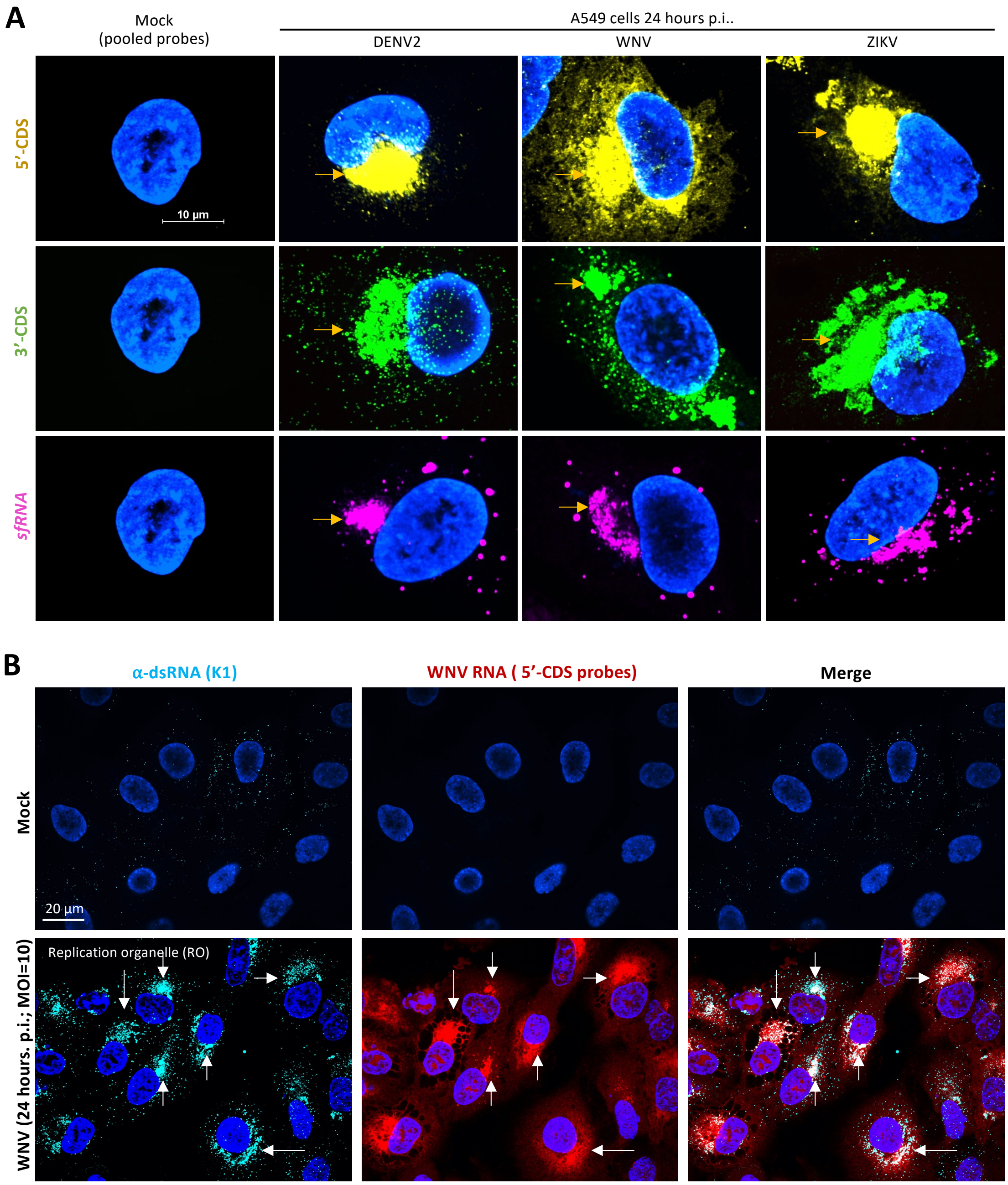


**Fig. S1. (A)** Individual staining examples of these probes during flavivirus infection. Yellow arrows correspond to replication organelles **(B)** IF for dsRNA co-stained with 5' WNV smFISH probes, showing replication organelle overlap. White arrows correspond to replication organelles.

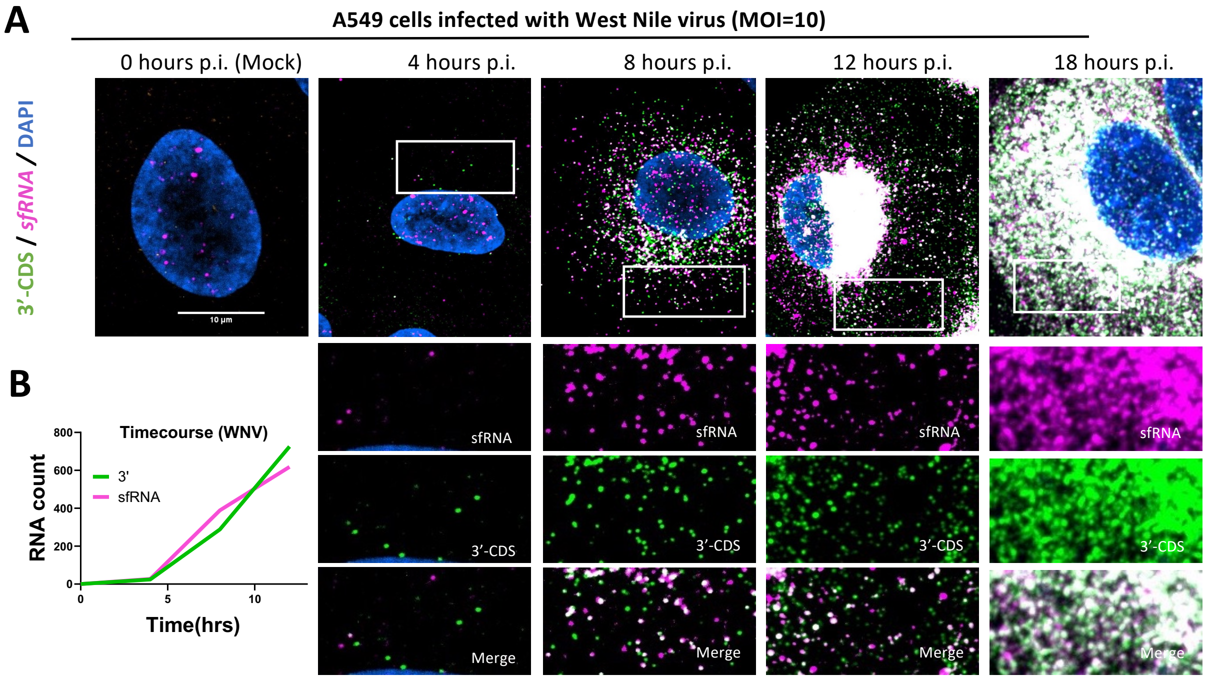


6
7 **Fig. S2. (A)** Representative images of 3'-CDS and sfRNA staining ranging from pre-infection to
8 18 hours p.i. with WNV **(B)** Quantitation of average RNA count per cell from (C).

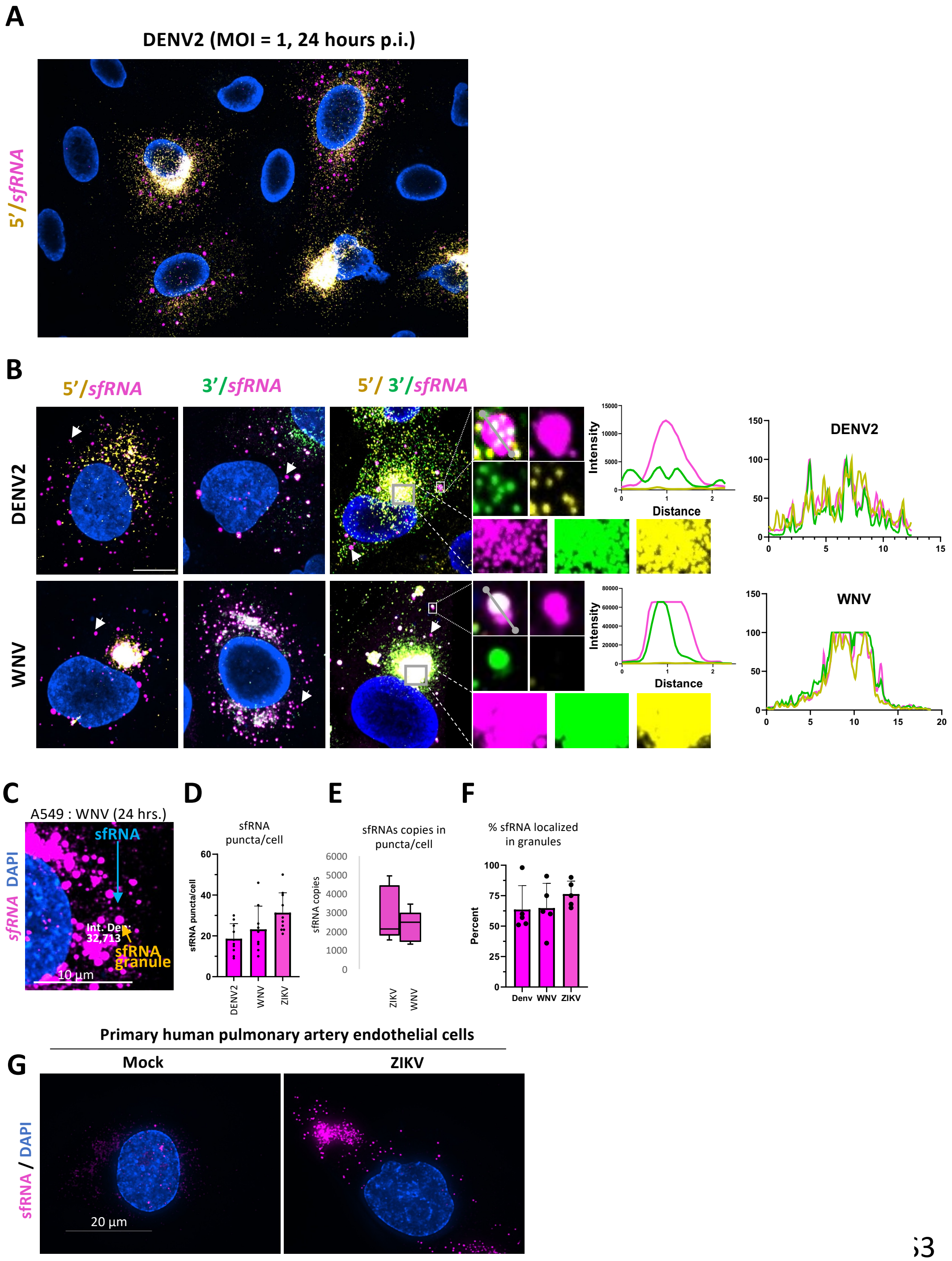


**Fig. S3. (A)** Representative image of infected cells co-stained for 5'-CDS and sfRNAs 24 hours p.i. with DENV2. **(B)** Co-staining permutations of DENV2 (MOI=0.1) and WNV (MOI=10), alongside line traces showing overlap in puncta favors 3' ends, while RO staining is equivalent. **(C)** Example image showing single sfRNA staining intensity vs inside a typical sfRNA puncta, 10um scale bar. **(D)** Quantitation of sfRNA puncta per infected cell 24 hours p.i. during DENV2 (MOI=0.1), WNV (MOI=10), and ZIKV (MOI=10) infections. **(E)** Number of sfRNA copies localized to puncta in individual cells, calculated using average intensity and number of puncta per cell. **(F)** Percent of total sfRNA localized in granules, as measured by fluorescent intensity. **(G)** Infection of HPAEC cells 24 hours p.i., showing sfRNAs colocalized with RLBs.

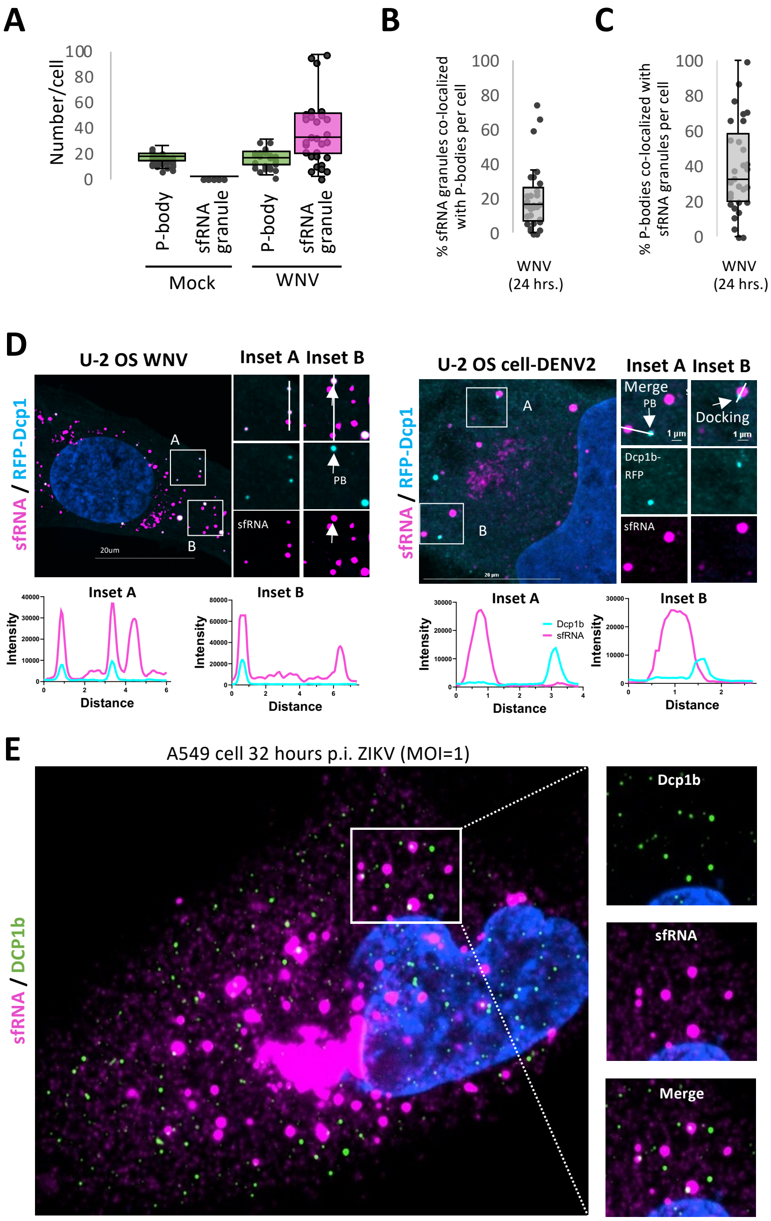


**Fig. S4. (A-C)** Quantitation of puncta as represented in (Fig. 2G). **(D)** Representative images of infection of U-2 OS cells containing RFP-Dcp1b, showing docking of sfRNA granules with P bodies in DENV2 (MOI=0.1) and WNV (MOI=10) infections 24 hours p.i. **(E)** Representative image of A549 cells infected with ZIKV co-stained with IF for DCP1b and smFISH for sfRNAs.

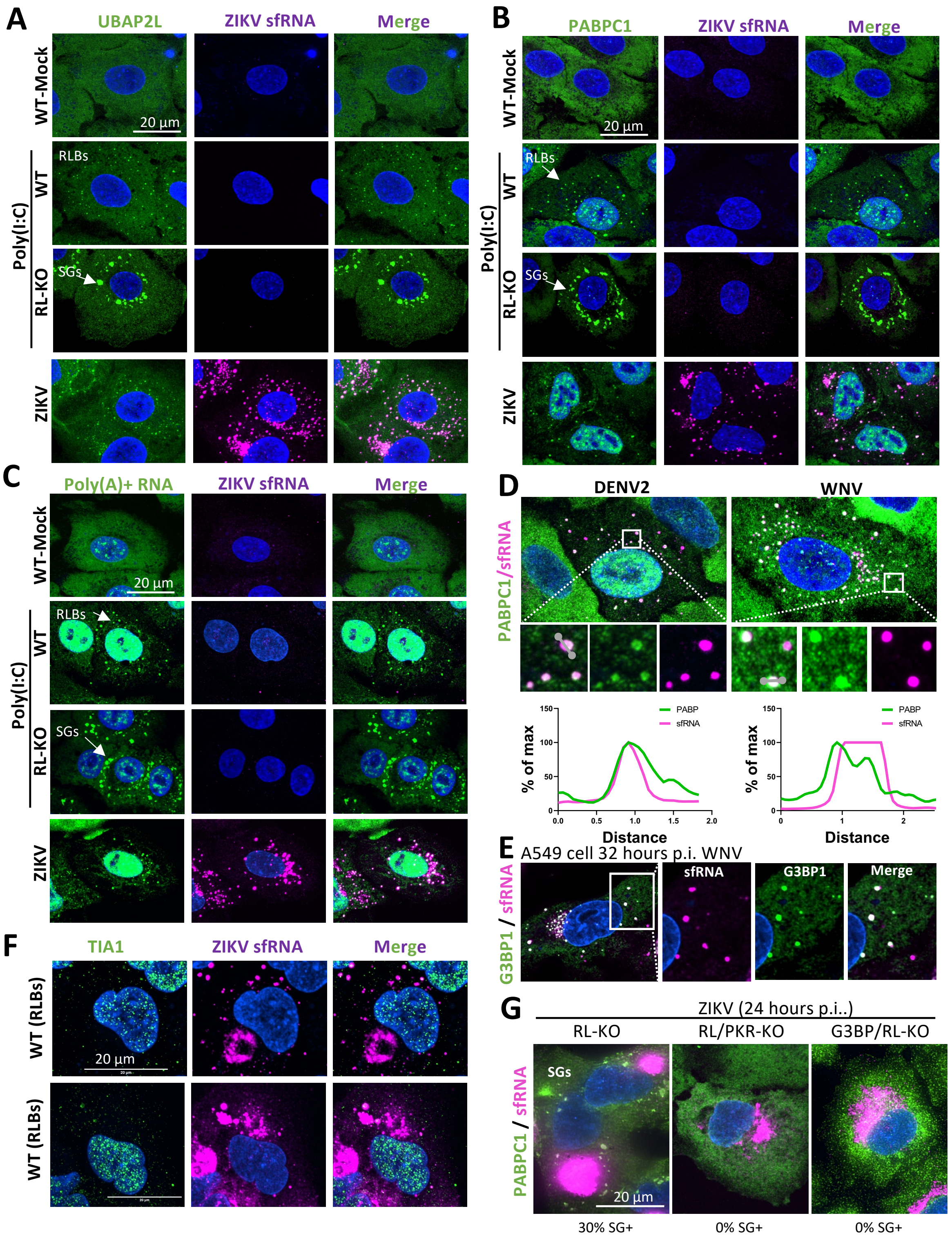


**Fig. S5. (A-C)** Representative images of IF for known RLB markers co-localizing with ZIKV (MOI=10) sfRNAs. Poly(I:C) images show RLBs (WT) or SGs (RL-KO cells). **(D-F)** Representative images showing DENV2 and WNV *sfRNA* smFISH with PABPC1 IF, G3BP1, and TIA1. **(G)** Co-staining of smFISH for sfRNA and IF for PABPC1 in various knockout cells.

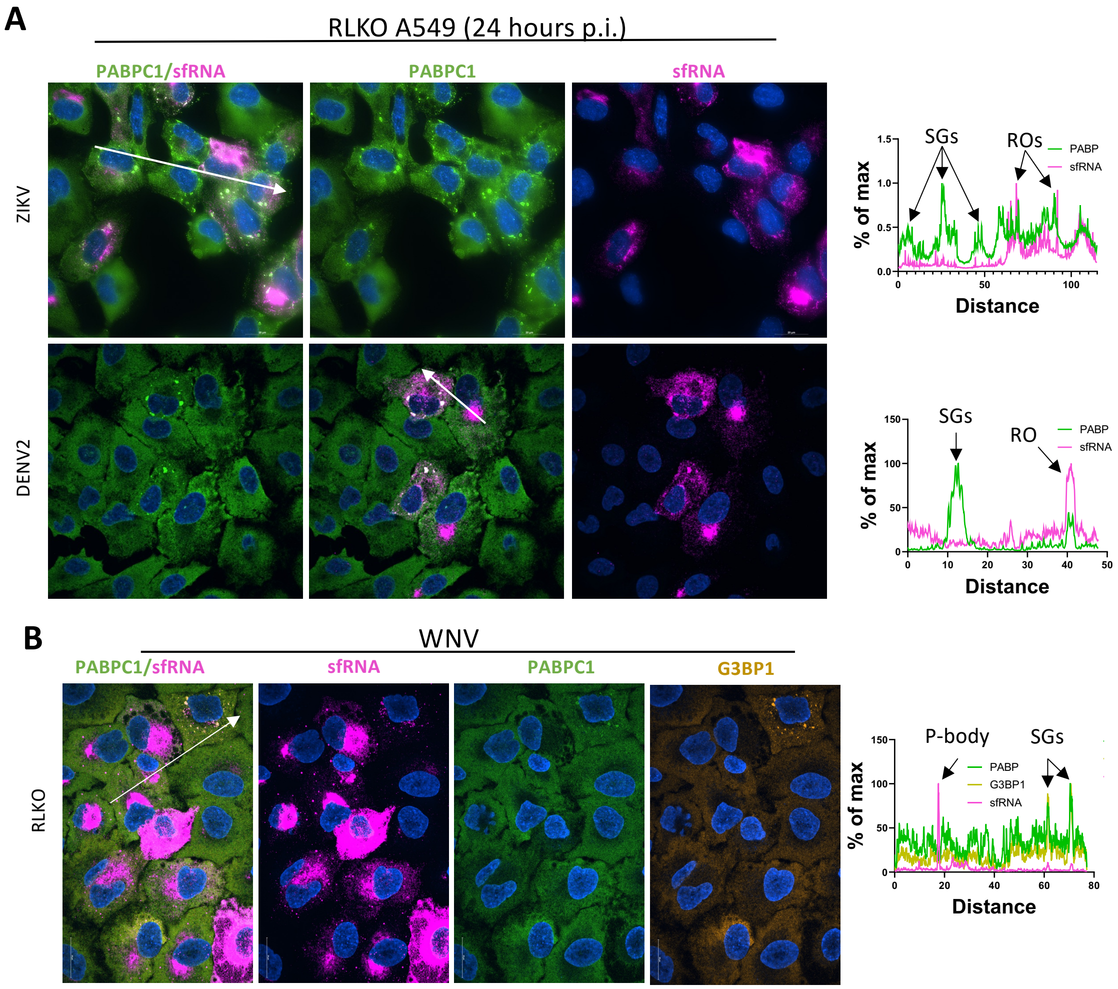


**Fig. S6. (A)** Representative wide-field images during ZIKV (MOI=10) and DENV2 (MOI=0.1) infection in RLKO A549 cells. IF of PABPC1 and *sfRNA* smFISH, with a line trace shown by white arrow. **(B)** Imaging of WNV (MOI=10) 24 hours p.i. Co-staining of PABPC1, G3BP1, and ZIKV (MOI=10) sfRNA. Line trace shows enrichment in P bodies vs in SGs.

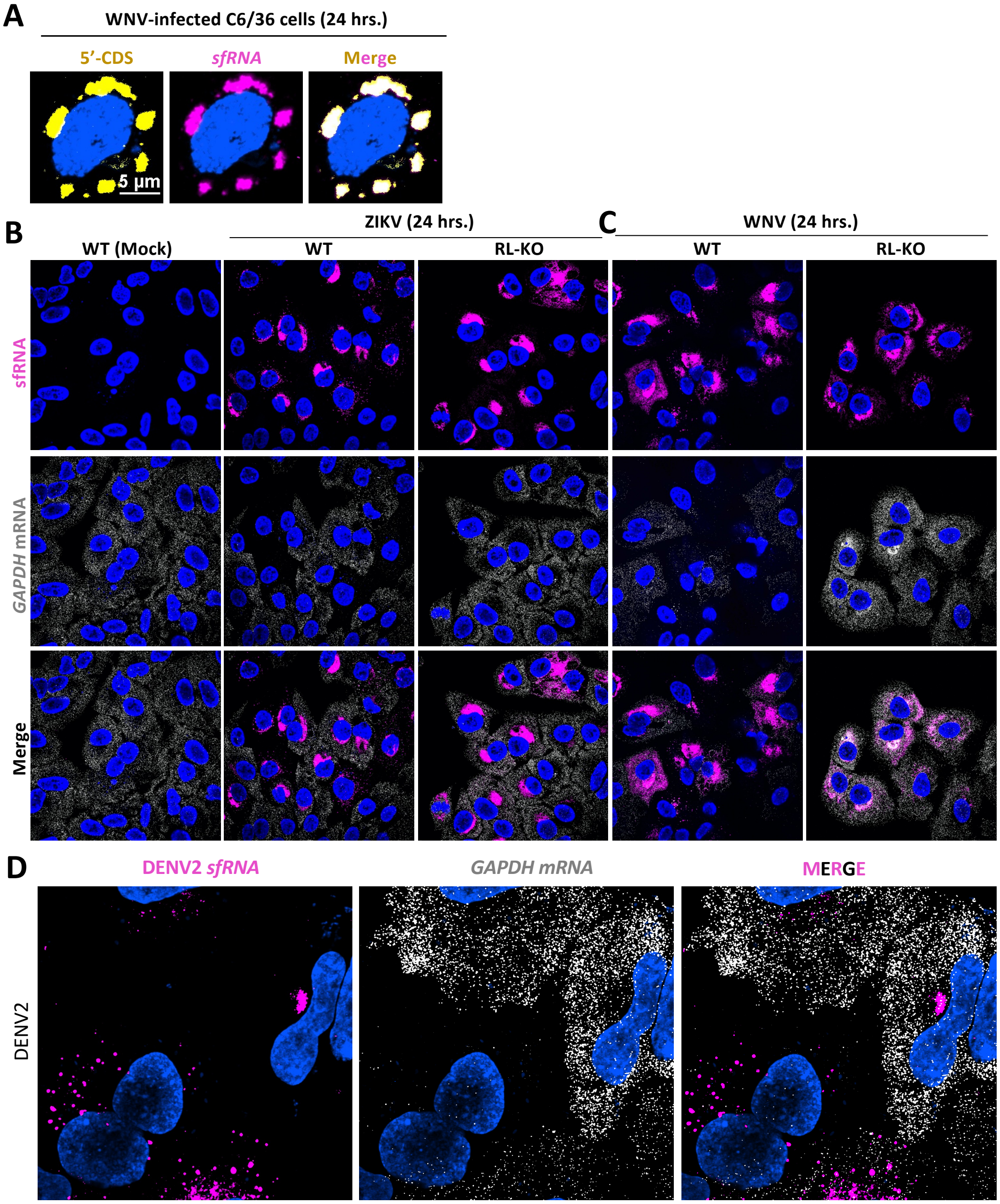


**Fig. S7. (A)** Representative imaging of insect cells, showing no independent sfRNA puncta formation in C6/36 cells. **(B-D)** Representative imaging of ZIKV (MOI=10), WNV (MOI=10), and DENV2 (MOI=0.1) infection, co-staining of *GAPDH* and *sfRNA* smFISH.

**A**

| | WT | RLKO |
|---|---|---|
| sfRNA1 | 1.00 | 0.90 |
| sfRNA2 | 1.00 | 0.12 |
| sfRNA3 | 1.00 | 0.43 |
| Total | 3.00 | 1.45 |

**B**

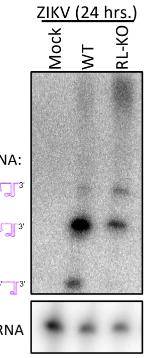


**Fig. S8.**

**(A)** Relative quantitation of intensities of each sfRNA in WT vs RLKO A549 cells during WNV (MOI=10) infection based on Northern blot analysis. Intensities were normalized to U6 RNA levels. Individual sfRNA bands were normalized to levels seen in WT A549 cells. **(B)** Northern blot analysis of ZIKV (MOI=10) sfRNA with U6 RNA load control.

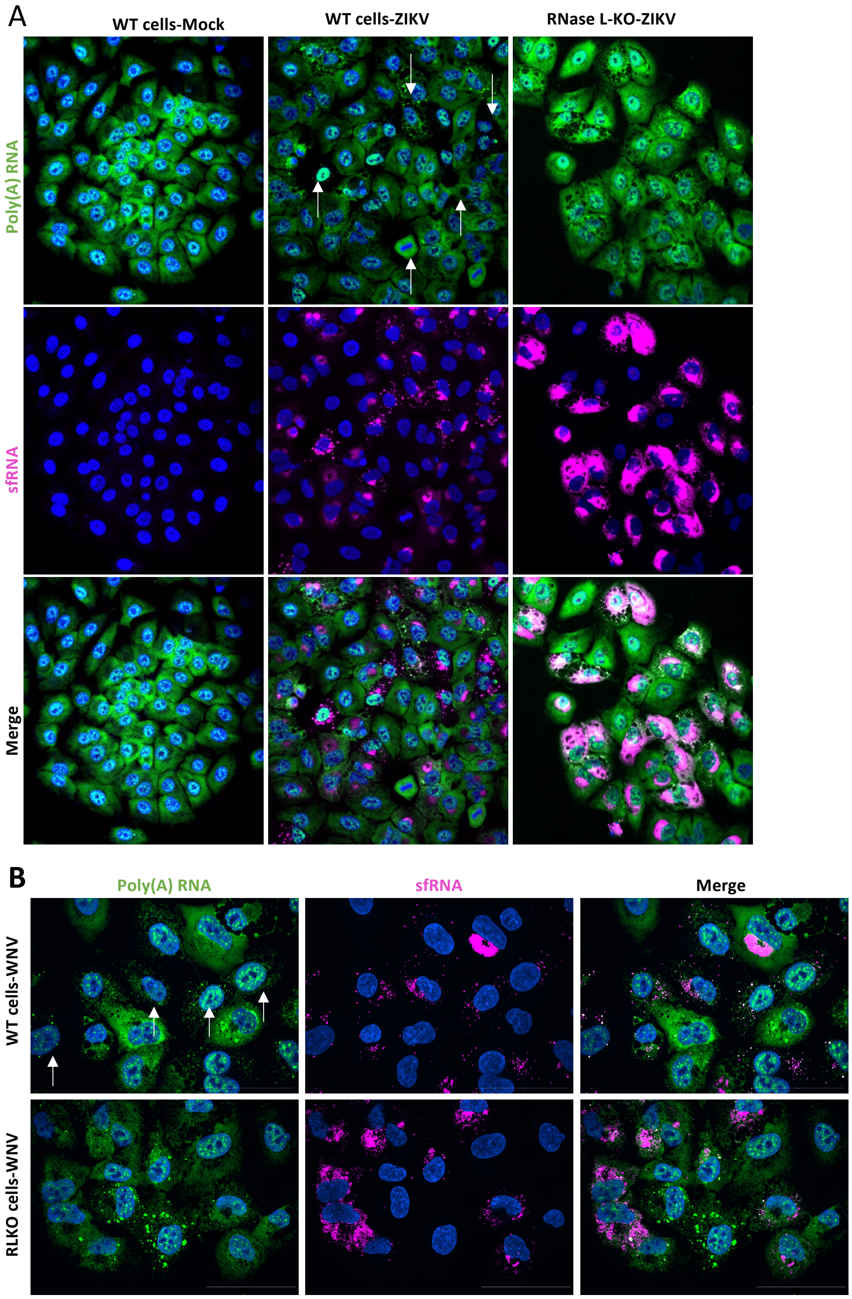


**Fig. S9. (A-B)** Wide-field imaging of FISH for poly(A) RNA and smFISH for sfRNA during ZIKV and WNV infection in WT and RLKO cells.

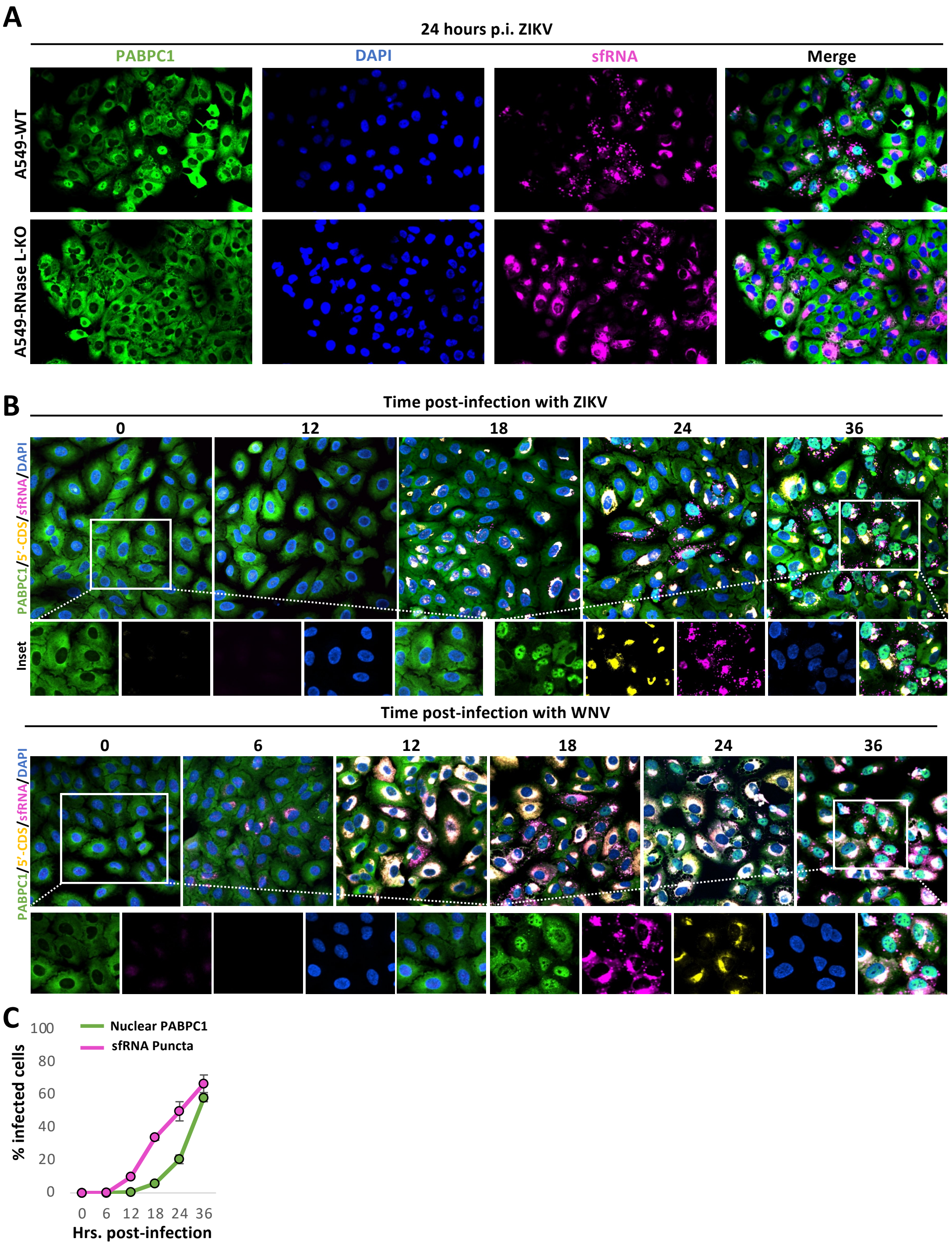


**Fig. S10. (A)** Wide field imaging of IF for PABPC1 and smFISH for sfRNA in WT and RLKO cells. **(B-C)** Wide field imaging of IF for PABPC1 and smFISH for 5'-CDS and sfRNA during ZIKV (MOI-=10) or WNV (MOI=10) timecourse. **(C)** Quantification of PABP translocation and sfRNA puncta formation during WNV infection.

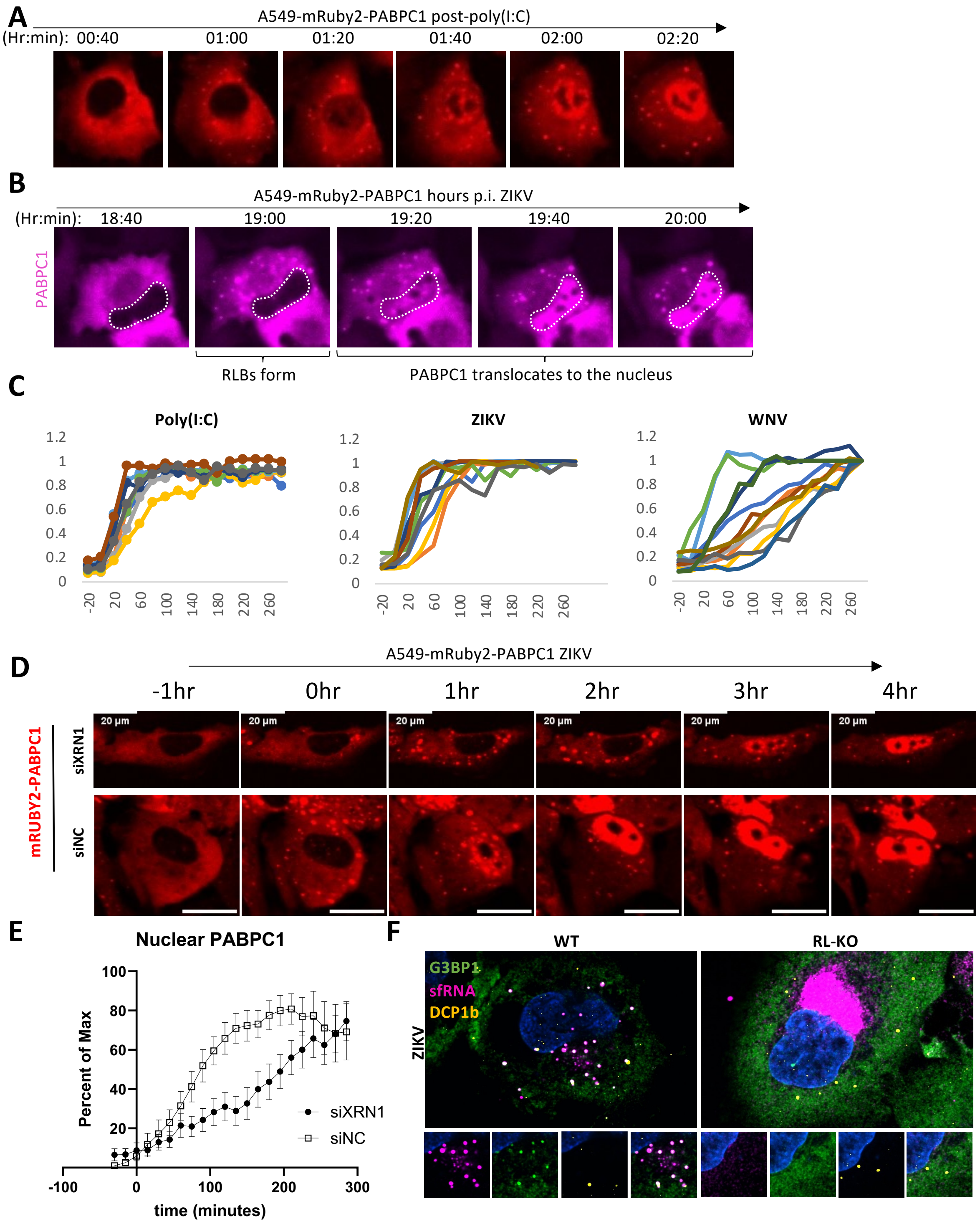


**Fig. S11. (A-B)** Live-cell imaging of mRuby2-PABPC1 translocation during p(I:C) and ZIKV infection. **(C)** Quantification of translocation rates of mRUBy2-PABPC1 during poly(I:C), ZIKV, and WNV infection. **(D)** Live-cell imaging of mRuby2-PABPC1 translocation during p(I:C) and ZIKV infection in cells with XRN1 or control knockdown. **(E)** Quantification 15 cells from (D). (**E**) smFISH for ZIKV sfRNA and IF for G3BP1 (SG/RLB marker) and DCP1B (P-body marker) in WT and RNase L-KO A549 cells 24 hours p.i.

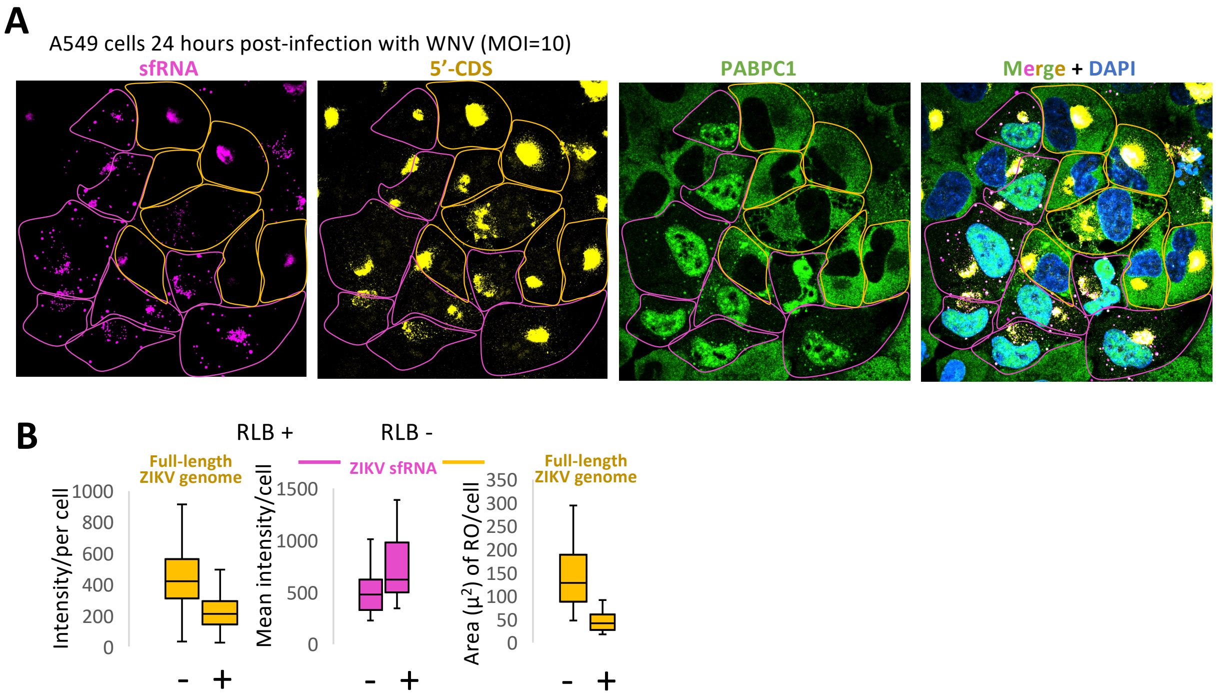


**Fig. S12. (A)** Co-staining of smFISH of sfRNA, 5'-CDS, and IF for PABPC1 24 hours p.i. Pink outlines indicate RNase L activation, while yellow outlines indicate inactive RNase L. **(B)** Quantification of intensity and size of replication organelle in cells with or without RNase L active from (A).
